# MsyB-HU Interaction Modulates a Two-Tiered Bacterial Stress Response by Regulating DNA Supercoiling

**DOI:** 10.1101/2025.02.25.640118

**Authors:** Huan Chen, Yanan Yang, Xinrui Chen, Haolong Zhou, Yingli He, Mengzhe Li, Wei Lin, Zuyi Yuan, Yawen Wang, Steve Matthews, Shuai Yuan, Hongliang Wang, Bing Liu

## Abstract

Bacteria respond to stressful conditions through multiple pathways, including changes in gene expression patterns and mutations for adaptation. DNA supercoiling is a fundamental regulatory principle of bacterial gene expression, controlled antagonistically by DNA gyrase and topoisomerase I. Nucleoid-associated proteins, such as HU, also regulate supercoiling through interactions with DNA. Here, we identify *Escherichia coli* protein MsyB as a HU inhibitor and a β-clamp binder. MsyB is fine-tuned under various stressful conditions, where it modulates the population of DNA-bound HU by forming a HU-MsyB complex. Consequently, the balance between DNA-bound HU and MsyB-bound HU adjusts the DNA supercoiling state, thereby influencing transcription for adaptation. Additionally, under prolonged starvation during the stationary phase, sustained high level of MsyB increases DNA exposure to damaging factors, acting as a damage inducer. The HU/MsyB/β-clamp interaction suggests a model in which MsyB coordinates with HU and the β-clamp to facilitate damage acquisition and error-prone DNA repair. Thus, MsyB-dependent supercoiling regulation represents a novel two-tiered bacterial stress response mechanism in gene expression and adaptive mutation. Our findings reveal a previously unrecognized regulatory mechanism of bacterial supercoiling, positioning HU as a promising antibiotic target.

## INTRODUCTION

Unrestricted growth condition is rarely encountered by bacteria in nature, as they predominantly exist in fluctuating stressful environments (*1, 2*). Bacteria can rapidly respond to stress through regulatory networks that modulate gene expression, enabling quick adaptation to challenging external conditions (*3*). As a result, bacterial gene expression patterns are altered compared to the optimal growth phase, with the upregulation of genes required for combating deleterious stresses and the downregulation of genes that are not involved. Stress-responsive transcriptional regulation can be achieved through various mechanisms, including pathways controlled by alternative σ factors, ppGpp and DNA supercoiling (*4–6*). Among these, DNA supercoiling is considered to be acting at or near the top of the regulatory hierarchy (*7*). It influences transcription by affecting both initiation and elongation, serving as a non-conventional regulator in both eukaryotes and bacteria (*8*). As a global transcriptional regulator in bacteria, supercoiling also plays a crucial role in adjusting gene expression to environmental changes (*9*). In bacteria, negative DNA supercoiling is modulated by the supercoiling activity of topoisomerase II (gyrase) and the relaxing activity of topoisomerase I (topo I) (*10, 11*). However, under many stress conditions, both gyrase and topo I are often downregulated at the transcript and protein levels (*12–14*). Despite their regulatory significance, no model is currently available to explain their role in supercoiling regulation in response to environmental stresses (*15*).

Nucleoid-associated proteins (NAPs) also have important roles in regulating supercoiling, by binding, bending and bridging DNA (*16*). Among *Escherichia coli* (*E. coli*) NAPs, HU can induce and constrain supercoiling by bending and coiling DNA (*17–19*), and crosstalk with topo I (*20, 21*). As one of the most abundant NAPs, HU binds to DNA non-specifically therefore scattered throughout the whole bacterial genome, in contrast to other sequence-specific NAPs (H-NS, Fis, and IHF) that localized in certain areas (*22–24*). The interplays between HU and topoisomerases have been shown to shape the landscape of supercoiling (*20*). It is therefore been proposed that HU may help gyrase to increase supercoiling, as well as assist topo I for relaxing the DNA (*20*). Importantly, HU-DNA interaction, likely through its role in adjusting DNA supercoiling, is considered as a general mechanism for bacterial transcriptional regulation during the cell cycle and stress responding (*25*). However, a regulatory mechanism that can effectively adjust HU activity also appears to be missing.

Some stress response pathways also allow bacteria to acquire mutations for adaptation (*26*). During stationary phase, for example, nondividing bacteria can also produce mutations in response to selective pressures through a process known as adaptive mutation (*27–31*). Adaptive mutation has an unusual high mutation rate (*32*), which is characterized by the involvement of DNA recombination/repair proteins and the error-prone DNA polymerase IV (Pol IV) (*33*). As the cell is hardly replicating or experiencing additional external stimuli that could induce DNA damage, the origin causing the rise of DNA damage is not clear. Notably, during stationary phase, the bacterial genome undergoes global compaction, facilitated by NAPs including HU, to protect DNA from damage (*34*), in contrast to the additional damage needed to increase mutation. Thus, the mechanism that removes these protections and introduces DNA damages remains largely unknown. Furthermore, even if DNA damages were induced, the DNA Pol IV does not seem to have a privileged access to damage sites to carry out repair (*35, 36*). As Pol IV should be granted the immediate access to the recombination intermediate site for the low-fidelity repair, a recruitment mechanism is needed to facilitate the entry of Pol IV. However, enigmatic mechanisms required to initiate these recombination-based mutations and to coordinate the damage induction and repair, remain unknown. To be noted, bacterial β subunit of DNA polymerase (β-clamp or clamp) is required for Pol IV to bind to DNA (*37*), and HU is shown to be involved in adaptive mutation via unknown mechanisms (*38*). Compared with the reversible stress response via transcription regulation, adaptive mutation is permanent and requires longer time to accumulate and therefore should occur no earlier than the change of transcription pattern during the bacterial stress-responding process.

Here, we report *E*. *coli* protein MsyB as a novel supercoiling regulatory protein for bacterial stress adaptation. MsyB levels are modulated in all tested stress conditions, with upregulation observed during stresses like starvation and antibiotic exposure, but downregulation during the SOS response. Since the HU-MsyB interaction is significantly stronger than the HU-DNA interaction, bacteria can regulate the amount of DNA-bound HU by adjusting MsyB expression. Consequently, the balance between DNA-bound HU and MsyB-bound HU allows bacteria to fine-tune DNA supercoiling, thereby modulating transcription for stress adaptation. Furthermore, by displacing large portion of total HU from DNA thus exposing DNA for damage acquisition, MsyB also functions as an inducer for DNA damage during adaptive mutation. Its interaction with the β-clamp, facilitated by a divergent clamp-binding sequence (F)QLEPP, enables the formation of the HU-MsyB-clamp complex. This suggests a model in which bacteria produce MsyB to induce DNA damage while coordinating β-clamp and Pol IV for low-fidelity repair, ultimately leading to mutations. Thus, by fine-tuning DNA-bound HU and interacting with the β-clamp, MsyB plays a critical role in two distinct bacterial stress responses: (i) transcriptional regulation for temporary adaptation and (ii) adaptive mutation at the genomic level for long-term survival.

## Results

### MsyB binds to β-clamp

Besides the ability to restore the growth and protein translocation defects in a temperature-sensitive *E. coli* mutant (*secY24*), little is known for the biological role of MsyB (*39*). However, one proteomic study shown that MsyB co-precipitated with β-clamp in a tandem affinity purification (TAP) experiment (*40*), yet absent in other similar studies (*41, 42*). To verify the interaction, we determined the affinity between MsyB and β-clamp at K_D_ = 0.70 μM using bio-layer interferometry (BLI, and Fig. 1A). As the stoichiometry of the interaction is yet to be determined, the result shows MsyB is a β-clamp binder but the interaction needs further characterization. We then employed the bacterial adenylate cyclase two-hybrid (BACTH) system to verify the interaction *in vivo* (Fig. 1B). The results agree with the BLI, showing interaction between MsyB and β-clamp in *E. coli,* which is native environment for this interaction. Interesting, MsyB only interacts with β-clamp when the adenylate cyclase tag is attached to its N-terminus, while β-clamp interacts with tagged MsyB regardless the position of the tag. Therefore, our data suggests that MsyB is another β-clamp interactor, which binds to β-clamp likely through its C-terminal region as a C-terminal tag disrupts their interaction. As MsyB does not contain the consensus β-clamp binding sequence QxxL(x)F (*43*) or QL(S/D)LF (*44*), which is likely the reason it was not previously identified as a β-clamp binder.

**Figure 1.**
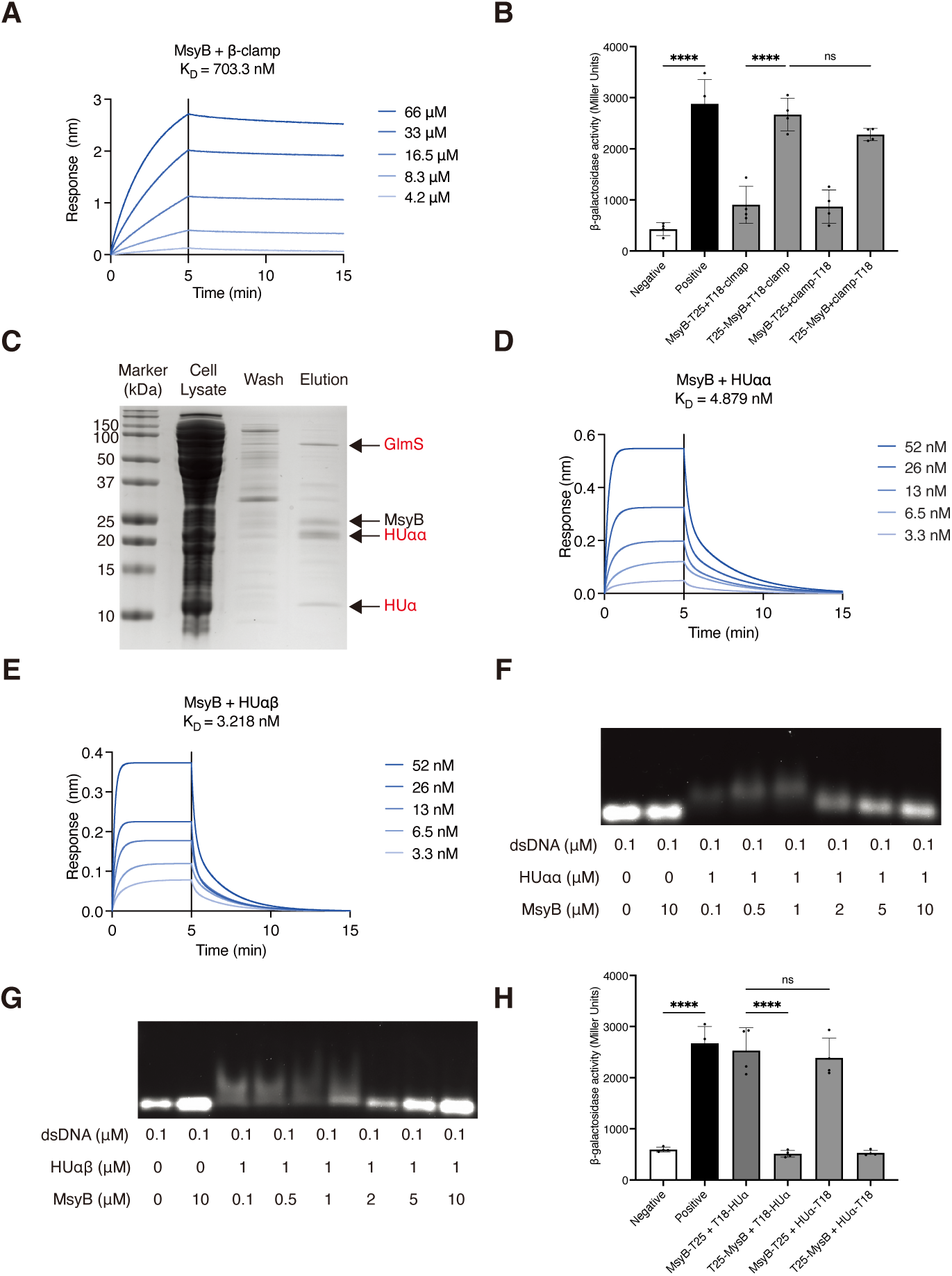
MsyB is a β-clamp interactor and HU inhibitor. **(A)** The affinity determined by BLI for the MsyB-clamp interaction. **(B)** The BACTH assays showing the interactions between MsyB and β-clamp *in vivo*, including interference from the positions of the tags. **(C)** SDS-PAGE showing the results of the pull-down experiment using column-immobilized MsyB with a C-terminal polyhistidine tag and cell lysate of *E. coli*. The bands at approximately 70 kDa and 10 kDa were determined as GlmS (a common contamination in polyhistidine-based purification) and HUα, respectively. The visible bands are labelled. **(D)** The affinity determined by BLI for MsyB-HUαα interaction. **(E)** The affinity determined by BLI for MsyB-HUαβ interaction. **(F)** The image of an agarose gel showing the ESMA assay demonstrating the competition between MsyB and dsDNA for HUαα. The final concentrations of HUαα and MsyB for each lane are shown in µM as indicated, with 0.1 µM dsDNA in all samples. The sequence of the dsDNA fragment is listed in supplementary table 1. **(G)** The image of an agarose gel showing the ESMA assay demonstrating the competition between MsyB and dsDNA for HUαβ, as demonstrated in (F). **(H)** The BACTH assays showing the interactions between MsyB and HUαα *in vivo*, including interference from the positions of the tags.

### MsyB is a native HU inhibitor

The β-clamp was first identified as the only MsyB interactor when an N-terminal tag was fused to the MsyB, thus the potential interactor that binds to the N-terminal region may be overlooked. To verify the speculation, we fused a C-terminal His-tag to MsyB and a SUMO tag to the N-terminal (SUMO-MsyB-His). We then column-immobilized MsyB-His after removing SUMO and performed a pull-down assay using the whole cell extract of *E. coli*.

We observed three major bands at approximately 10 kDa, 22 kDa and 65 kDa that co-eluted with MsyB in the SDS gel (Fig. 1C). The corresponding proteins which were then identified as the HUα, HUαα (dimer) and Glutamine--fructose-6-phosphate aminotransferase (GlmS) by mass spectrometry (MS), respectively (Fig. S1A-C). As GlmS is a common contamination in polyhistidine-based purification, HU is likely the primary interactor of MsyB. The BLI determined the K_D_ of the interaction to be 4.879 nM between HUαα and MsyB, and 3.218 nM between HUαβ and MsyB (Fig. 1D and E). Importantly, the affinities between HU and MsyB are much stronger than that between HU and double-stranded DNA (dsDNA) (293 nM) (*45*), but they are comparable to the affinities between HU and its preferred DNA substrates, including junctions and nicks, which ranging from 1.6 to 16 nM (*46*). Considering its potential DNA mimicry due to its very low pI (3.6), we also tested other major NAPs, including IHF, FIS, and H-NS, to determine if they interact with MsyB. The results show that MsyB interacts with IHF, a close structural homologue of HU, which binds to specific DNA sequences and sharply bends the DNA, but not with FIS or H-NS (Fig. S2A and B). The calculated affinity between IHF and MsyB (95 nM, Fig. S2C) is lower than between HU and MsyB, and the affinity between IHF and its cognate DNA sequence (0.5-20 nM) (*47*). Therefore, MsyB would prefer to interact with HU on plain dsDNA than to displace IHF from the consensus sites.

The high affinity for HU-MsyB interaction implies that MsyB is likely a HU inhibitor. An electrophoretic mobility shift assay (EMSA) confirmed that MsyB could displace DNA from the HUαα-dsDNA complex (Fig. 1F). Similar results were obtained for MsyB and HUαβ, confirming its ability to replace dsDNA from both biological relevant HU dimers (Fig. 1G). In the BACTH assay for verifying the HUαα-MsyB interaction, we attached a T25 fragment to either terminus of MsyB and a T18 fragment to either terminus of HUαα. The results indicated that only C-terminal tagged MsyB was able to interact with HUαα, while the position of the tag on HUα had no effect on the interaction (Fig. 1H).

### MsyB inhibits DNA supercoiling

*E. coli* contains 30,000 to 55,000 HU monomers, or approximately 25,000 HU dimers per cell as determined by quantitative western blot analysis (*48*). The population remains relatively constant during exponential phase but dropped to 15, 000 HU per cell during early stationary phase. Meanwhile, a quantitative and condition-dependent *E. coli* proteomic study revealed that MsyB has its smallest population of 1,792 molecules per cell during the exponential phase in LB medium, but its abundance increases at different scales under various stressful conditions, including nutrient-limitation, acidic stress, temperature and starvation (*14*) (Fig. S3A), which also is supported by the elevated levels of MsyB transcripts under different stresses (*12*). For example, the population of MsyB in one *E. coli* cell increases sixfold from exponential phase to stationary phase, while HU level drops more than 50%. Using RT-qPCR, we show that MsyB transcript levels increased more than 20-fold after entering stationary phase, supporting that finding of the proteomic study (Fig. S3B). Based on these numbers, if MsyB has a similar *in vivo* effect against HU, it will remove over majority of HU from DNA during stationary phase, compared to only a small fraction of HU during the exponential phase (assuming 1:1 ratio). Removing large amount of HU from DNA would significantly alter DNA supercoiling, therefore, theoretically, the MG1655^Δ*msyB*^ strain (Δ*msyB*), with all HU bound to DNA, and the MsyB-overexpressing MG1655 strain (MsyB^OE^), with most HU removed from DNA, should have distinguishable supercoiling profiles compared to the wild-type MG1655 strain (WT), whose HU exists in both DNA-bound and MsyB-bound forms.

To verified if the supercoiling is affected as anticipated, we used biotinylated trimethyl psoralen (bTMP), which preferentially binds to negatively coiled DNA and has been widely used in supercoiling profiling in eukaryotic cells (*49, 50*), to probe the DNA supercoiling states of *E. coli* during exponential and stationary phases. During the exponential phase, the normalized fluorescence intensities (AU per cell), are 402.7, 852.5, 520.9 and 2339.8 per cell for MsyB^OE^, MG1655 (WT), Δ*hupA/*Δ*hupB* (Δ*hupAB*) and Δ*msyB* strains, respectively (Fig. 2A and Fig. S3D). The bTMP-based supercoiling profiles of MsyB^OE^ and Δ*hupAB* strains were similar, which agrees with previous observations that deleting HU leads to a decrease in global negative superhelicity during both the exponential and stationary phases (*51, 52*), supporting that MsyB acts as a negative regulator of supercoiling via its interaction with HU. Moreover, the even lower normalized fluorescence intensity observed in MsyB^OE^ cells compared to Δ*hupAB* cells supports the notion that an excess amount of MsyB can also function as an IHF inhibitor, which is known as a “supercoiling buffer” (*53*). The inverse relationship between DNA supercoiling and MsyB expression levels suggests that MsyB inhibits supercoiling. Consistent with the decrease in average chromosomal supercoiling during stationary phase (*54, 55*), the fluorescence intensity for WT dropped from 852.5 to 475.4, while the overall supercoiling trend remained similar to that of the exponential phase (Fig. 2B and S3E). The fluorescence intensity of MsyB^OE^ strain was very close to that of WT in the stationary phase, which aligns with the significant upregulation of *msyB* during this phase.

**Figure 2.**
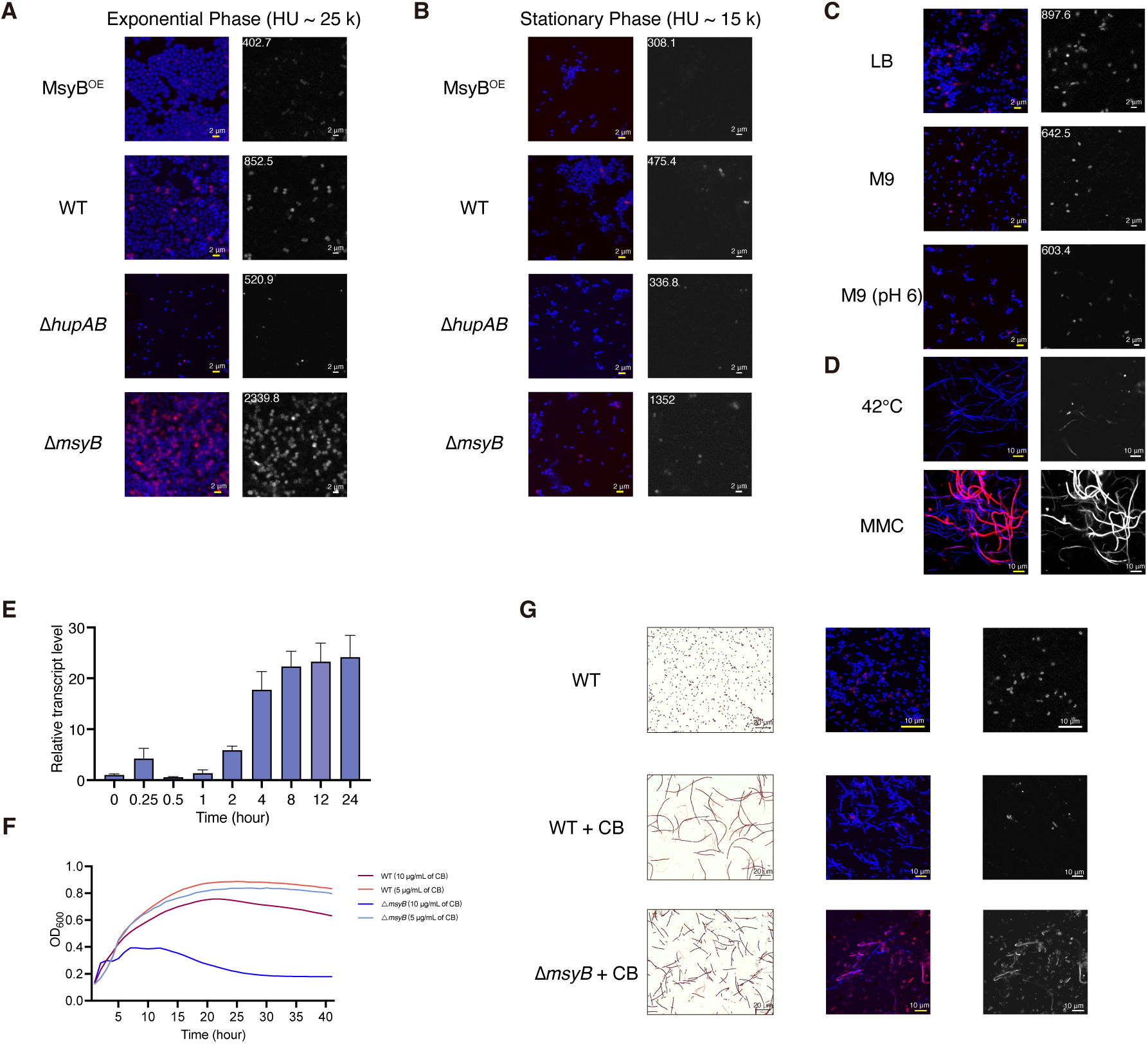
MsyB modulates DNA supercoiling. (**A-B**) The representative bTMP staining images of MsyB^OE^, WT, Δ*hupAB* and Δ*msyB* cells of exponential phase (**A**) or stationary phase (**B**) (right panels). Nucleoids were counterstained with DAPI and merged images were shown (left panels). Bar, 2 μm. The number represents the normalized florescence intensity of current frame. (**C**) The representative bTMP staining images of WT cells cultured in LB medium, M9 medium or M9 medium (pH 6.0) for 6 hours before they were stained with bTMP (right panels). Nucleoids were counterstained with DAPI and merged images were shown on the left. Bar, 2 μm. The number represents the normalized florescence intensity of current frame. (**D**) The representative bTMP staining images of WT cells cultured in M9 medium for 1 hour at 42°C or treated with 1μg/mL MMC for 6 hours at 37°C before they were stained with bTMP (right panels). Merged images were shown on the left. Bar, 10 μm. The number represents the normalized florescence intensity of current frame. (**E**) WT cells were treated with 10 μg/mL CB for the indicated time and the relative expression of *msyB* was determined by RT-qPCR. (**F**) WT or Δ*msyB* cells were treated with 5 or 10 μg/mL CB and the growth curved were measured. Three biological replicates were performed, each containing three technical replicates. The growth curve represents the average OD_600_ from a representative experiment. (**G**) WT or Δ*msyB* cells were treated with 10 μg/mL CB before they were Gram stained (left panels) or stained with bTMP (right panels). Nucleoids were counterstained with DAPI and merged images were shown in the middle. Bar, 10 μm.

### MsyB modulates DNA supercoiling under various stressful conditions

We then applied different challenges to *E. coli* and examined the transcript levels of *msyB* and supercoiling. Since the population of MsyB increases during nutrient limitation, acidic pH, and 42°C, we first verified this at the transcript level using RT-qPCR. The results agree with the proteomic data, showing significant upregulation of *msyB* under these stressful conditions (Fig. S4A). Meanwhile, MMC has the opposite effect on *msyB* levels; adding MMC to WT strain during the stationary phase in LB media represses *msyB* at the transcript level and flattens its expression during the exponential phase (Fig. S4B). The supercoiling profiles of *E. coli* under these conditions are inversely related to *msyB* levels, showing reduced supercoiling under nutrient limitation, acidic pH, and 42°C, but increased supercoiling under MMC treatment (Fig. 2C, 2D and S4C). However, when WT cells were cultured in M9 medium at 42°C or treated with MMC, they exhibited an elongated filamentous morphology, resulting in significantly larger and highly variable cell sizes compared to normal cells (Fig. 2D). Therefore, normalized fluorescence intensity was not used for quantification. In contrast to MMC-treated cells, which showed increased supercoiling, cells in M9 medium at 42°C exhibited lower supercoiling. And the growth curves indicated that the growth of the Δ*msyB* strain is not affected in LB but is impaired when exposed to these conditions (Fig. S4D), supporting that a tuned supercoiling profile is required for the optimal adaptation to specific environmental stress.

Additionally, we show that the transcript levels of *msyB* were upregulated in distinct patterns during exposure to sublethal concentrations of antibiotics, using carbenicillin (CB), tetracycline (TET), and levofloxacin (LFX) to represent three major antibiotic classes based on their mechanisms of action (Fig. 2E and S5A). The Δ*msyB* strain exhibited significantly impaired growth under antibiotic treatment compared to the WT (Fig. 2F and S5B). Using CB as an example, the phenotypes of CB-treated *E. coli* were intriguing: the WT cells exhibited the characteristic elongated shape under CB treatment, while most drug-treated mutant cells were significantly shorter and stained poorly (Fig. S5C). Subsequent methylene blue staining revealed that the shorter mutant cells were dead, whereas the longer ones remained alive (Fig. S5C). The remaining mutant cells were longer than the shorter mutants but still shorter than the WT cells. These observations suggest that the Δ*msyB* strain is unable to properly adjust its cell length, unlike the WT strain, to counteract the impaired cell wall synthesis caused by CB. The supercoiling profile also suggests that the Δ*msyB* cells still maintained a high level of supercoiling compared to the relaxed level seen in the WT cells (Fig. 2G). Together, these results demonstrate that supercoiling levels are modulated by the population of MsyB during these stressful conditions.

### MsyB-dependent supercoiling affects global gene expression during stressful conditions

To demonstrate that supercoiling affected by the absence of MsyB can lead to changes in gene transcription, we analyzed gene expression during starvation and antibiotic exposure using RNA-seq. Differential expression analysis between the stationary and exponential phases in WT cells revealed that 1,178 genes were upregulated, while 1,219 genes were downregulated during the stationary phase (Fig. 3A, upper panel and Table S4). In contrast, Δ*msyB* cells exhibited a smaller difference, with 957 genes upregulated and 971 genes downregulated (Fig. 3A, lower panel and Table S5). These results indicate reduced adjustment of gene expression in Δ*msyB* cells, similar to previous findings in Δ*hupAB* cells (*25*). Furthermore, the KEGG enrichment analysis shows significant difference in the enriched pathways (Fig. 3B). In WT cells, the biosynthesis of secondary metabolites, auxiliary compounds that are not essential for normal cell growth but are crucial for environmental adaptation (*56*), was significantly enriched. However, this enrichment was absent in Δ*msyB* cells.

**Figure 3.**
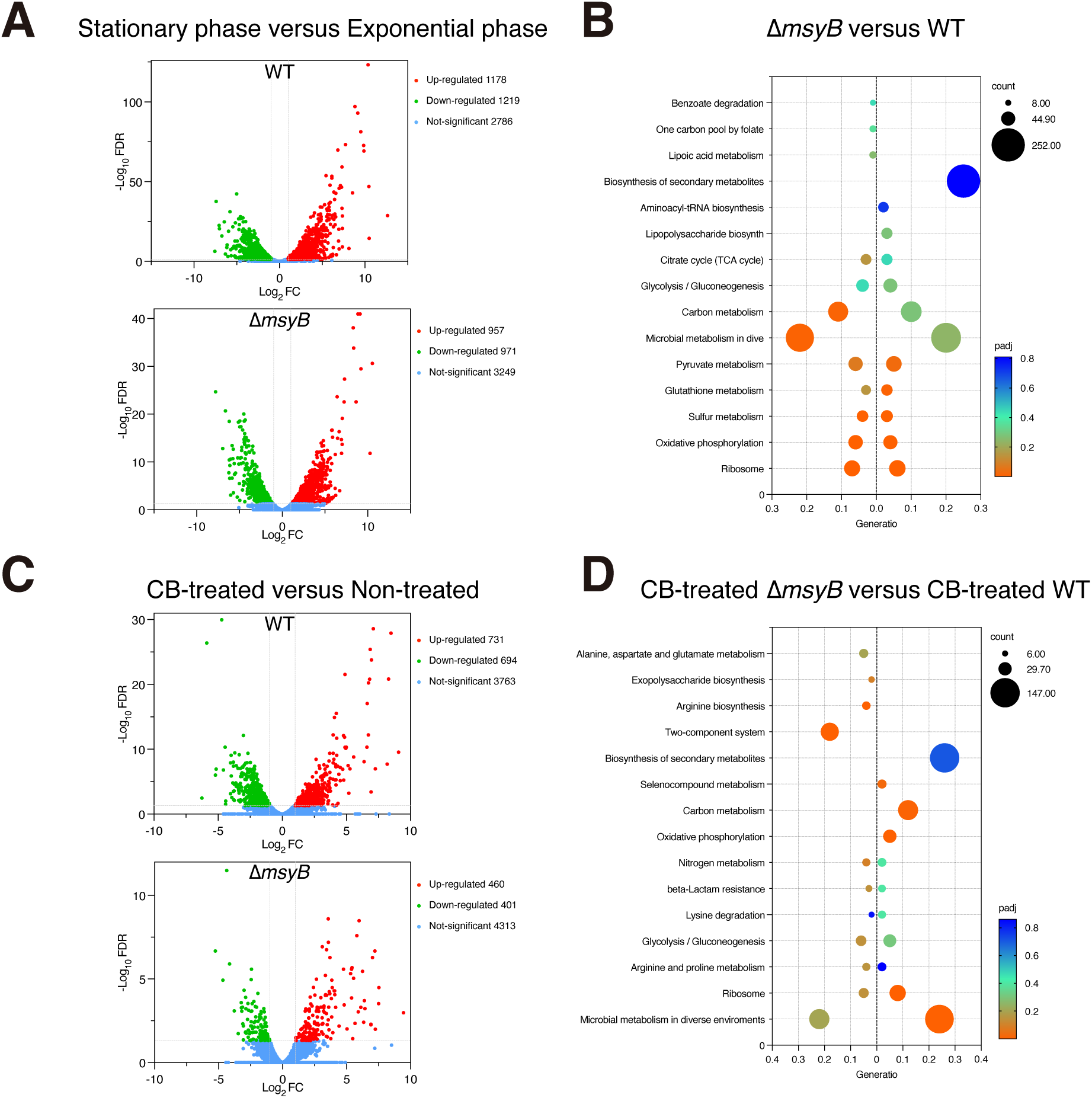
Global gene expression of WT and *ΔmsyB* cells under starvation and CB treatment. (**A&C**) Volcano plot showing log_2_ fold change (log_2_ FC) in transcript abundance plotted against -log_10_ false discovery rate (-log10 FDR). Two vertical bars indicate log_2_ FC cutoff of -1 and 1, while a horizontal bar represents an FDR cutoff of 1.301. Each point represents a gene; red points indicate upregulated genes, green points indicate downregulated genes, and blue points indicate unchanged genes. (A) Differential gene expression between the exponential and stationary phases in WT (upper) and Δ*msyB* (lower) strains, respectively. (**C**) Differential gene expression in WT (upper) and Δ*msyB* (lower) strains with and without CB treatment. **(B&D)** KEGG pathway enrichment analysis. Y-axis indicates the pathway name; x-axis indicates gene ratio. The bubble size indicates the number of genes and the color bar indicates the adjusted p-value. (B) Pathway enrichment of differentially expressed genes during bacterial growth from exponential phase to stationary phase in WT (right) and Δ*msyB* (left) strains, respectively. (D) Pathway enrichment of differentially expressed genes between CB treated and untreated cells in WT (right) and Δ*msyB* (left) strains, respectively.

Under the CB exposure, we observed that 731 genes were upregulated and 694 genes were downregulated in WT cells (Fig. 3C, upper panel and Table S6). In contrast, only 460 genes were upregulated and 401 genes were downregulated in Δ*msyB* cells (Fig. 3C, lower panel and Table S7). Similarly, KEGG analysis showed that the metabolism in diverse environments was the most enriched pathway in WT cells but was absent in Δ*msyB* cells (Fig. 3D), and showing that response of WT cells to the antibiotic stress. However, the two-component system, which enables bacteria to sense and respond to environmental changes (*57*), became enriched in Δ*msyB* cells, possibly playing a compensatory role due to the absence of MsyB. These results show that while bacteria can partially compensate for the loss of supercoiling regulation through other stress-response pathways, this compensation appears to be incomplete. These results support that the supercoiling level modulated by MsyB is a master regulator of global gene expression in response to starvation and antibiotic exposure.

### Cryo-EM study of HUαα-MsyB-clamp ternary complex

To elucidate the molecular basis of how MsyB regulates supercoiling through its interaction with HU and β-clamp, we employed structural tools to characterize these interactions. As BACTH suggests different parts of MsyB interact with β-clamp and HUαα, the MsyB-clamp and HUαα-MsyB interactions may be simultaneous. To verify the speculation, we first assembled HUαα-MsyB complex at 1:1 ratio (Fig. 4A, HU has no absorbance at 280 nm). And adding excess amount of HUαα to MsyB does not shift the peak, suggesting that the stoichiometry in complex is 1:1. Adding purified HUαα-MsyB complex to β-clamp results in a migration in ÄKTA, suggesting the formation of a larger complex than β-clamp alone. As excess amount of HUαα-MsyB does not change the elution volume of the putative HUαα-MsyB-clamp complex, the stoichiometry of the HUαα-MsyB-clamp complex is therefore1:1:1 (Fig. 4B). The affinity between HUαα-MsyB and β-clamp was determined using HUαα-MsyB complex and β-clamp at K_D_ = 499.5 nM by BLI (Fig. 4C). The higher affinity suggests that HU enhances rather than disrupts the interaction between MsyB and β-clamp. And we then assembled the HUαα-MsyB-clamp ternary complex for cryo-electron microscopy (cryo-EM) single particle structure determination.

**Figure 4.**
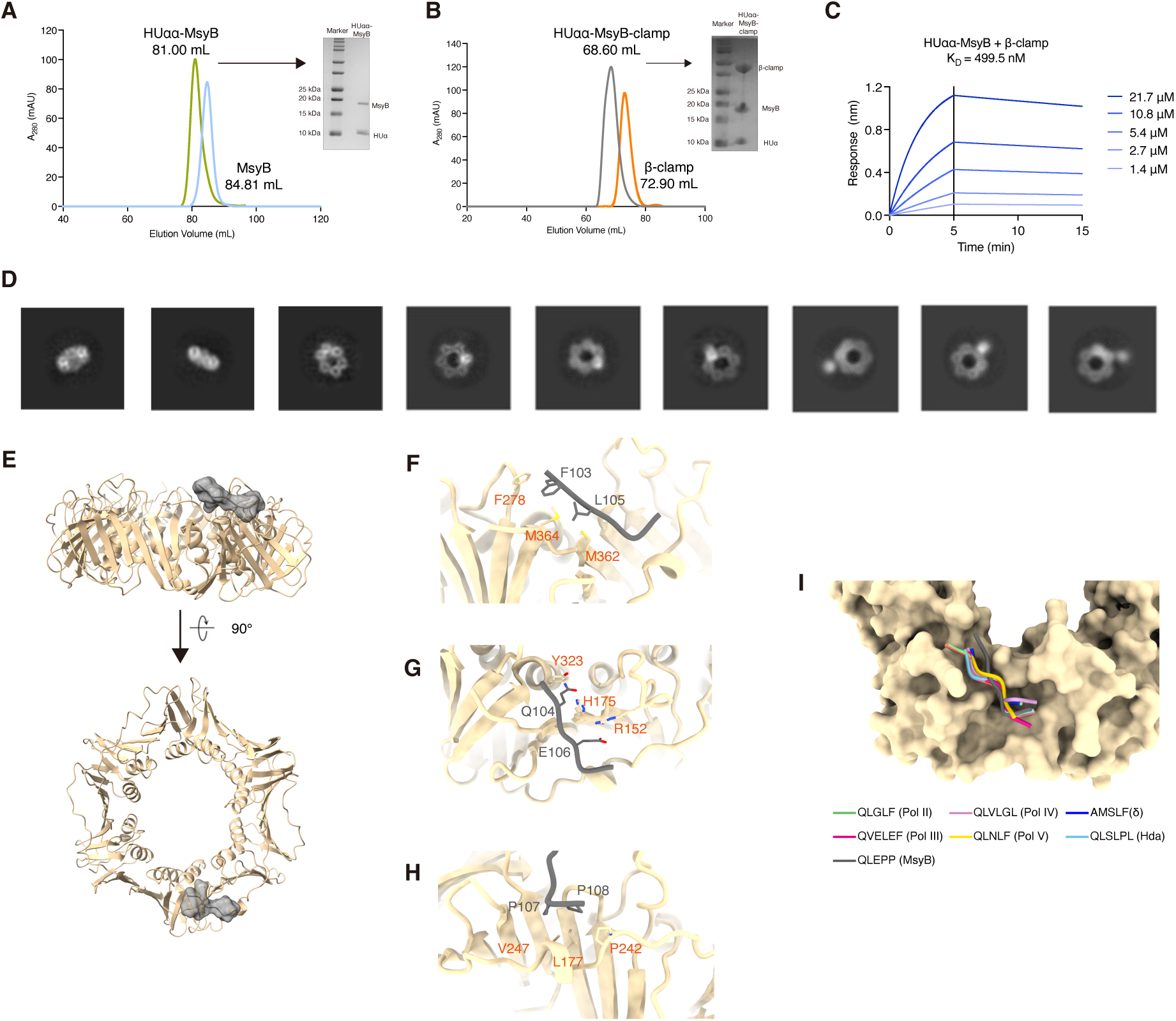
Cryo-EM studies of HU-MsyB-clamp complex. **(A)** The gel filtration profiles of MsyB, and HU-MsyB (1:1) complex assembled by adding equimolar MsyB to HUαα. And the accompanied SDS-PAGE showing the formation of the HU-MsyB complex. **(B)** The gel filtration profile of clamp and HU-MsyB-clamp (1:1:1) complex assembled by adding equimolar HU-MsyB to clamp. And the accompanied SDS-PAGE showing the formation of the HU-MsyB-clamp complex. **(C)** The affinity determined by BLI for the HU-MsyB and β-clamp interaction. **(D)** 2D classification of HU-MsyB-clamp complex **(E)** The cryo-EM structure of MsyB-clamp (fragment: QLEPP) complex. **(F-H)** The interactions between residues of β-clamp and MsyB. **(I)** The superimposing of the fragments from different β-clamp interactors (Pol II, III, IV, V, δ and Hda) in the protein binding pocket of the β-clamp.

In the 2D classification, we observed extra densities at variable distances from the centre of the clamp ring, presumably the HUαα-MsyB binding to only one of the two protein binding pockets (Fig. 4D and Fig. S6A). The observation could be explanation by the potential existence of C-terminal intrinsically disordered (ID) domain in MsyB based on the disorder prediction using IUPred3 (*58*). Although this flexibility complicates the structural determination of the complex, we managed to obtain a complex containing a fragment of MsyB (QLEPP) that binds to the clamp in one of the protein-binding pockets (Fig. 4E, S6B and C). The vacant pocket on the clamp ring also confirms the 1: 1: 1 stoichiometry of the complex. The interactions between MsyB and β-clamp can be categorized into three sites: F103 and L105 of MsyB interact with F278, M362 and M364 of β-clamp through hydrophobic interactions (Fig. 4F); Q104 of MsyB forms hydrogen bonds with Y323 and H175 of β-clamp, and E106 of MsyB interacts with R152 of β-clamp via electrostatic interaction (Fig. 4G); and P107 and P108 of MsyB engage with L177, V247, and P242 of β-clamp through hydrophobic interactions (Fig. 4H). As known *E. coli* clamp-binding proteins contain a conserved sequence QxxL(x)F or QL(S/D)LF that binds to the hydrophobic pocket on the clamp ring, the divergent QLEPP sequence in MsyB binds to the hydrophobic pocket in a similar manner (Fig. 4I), albeit with variations compared with other known binders (*59*). In summary, the cryo-EM studies reveal the flexibility within the HUαα-MsyB-clamp ternary complex and show that MsyB contains a divergent β-clamp binding sequence, occupying only one of the two hydrophobic pockets on the clamp ring.

### HUαα-MsyB-clamp tripartite complex model

To complement the EM studies, we determined the solution structure of MsyB by using standard multidimensional NMR experiments (*60*) (Table S2 and PDB ID: 8IMQ). Notably, the solution structure contains two distinctive domains: an N-terminal globular domain and a C-terminal intrinsically disordered region (IDR) (Fig. 5A). The N-terminal domain (NTD) folds into a globular structure that does not resemble any published structure as homology search using Dali server (*61*) per suggested. The lack of identifiable structure in the C-terminal domain (CTD) is consistent with the disorder predictions (Fig. S7A). Using the structure closest to the average solution structure from the NMR ensemble, we calculated the electrostatic surface of MsyB (Fig. S7B). Unsurprisingly, the surfaces of both NTD and CTD are predominately occupied by negative charge as MsyB is very acidic.

**Figure 5.**
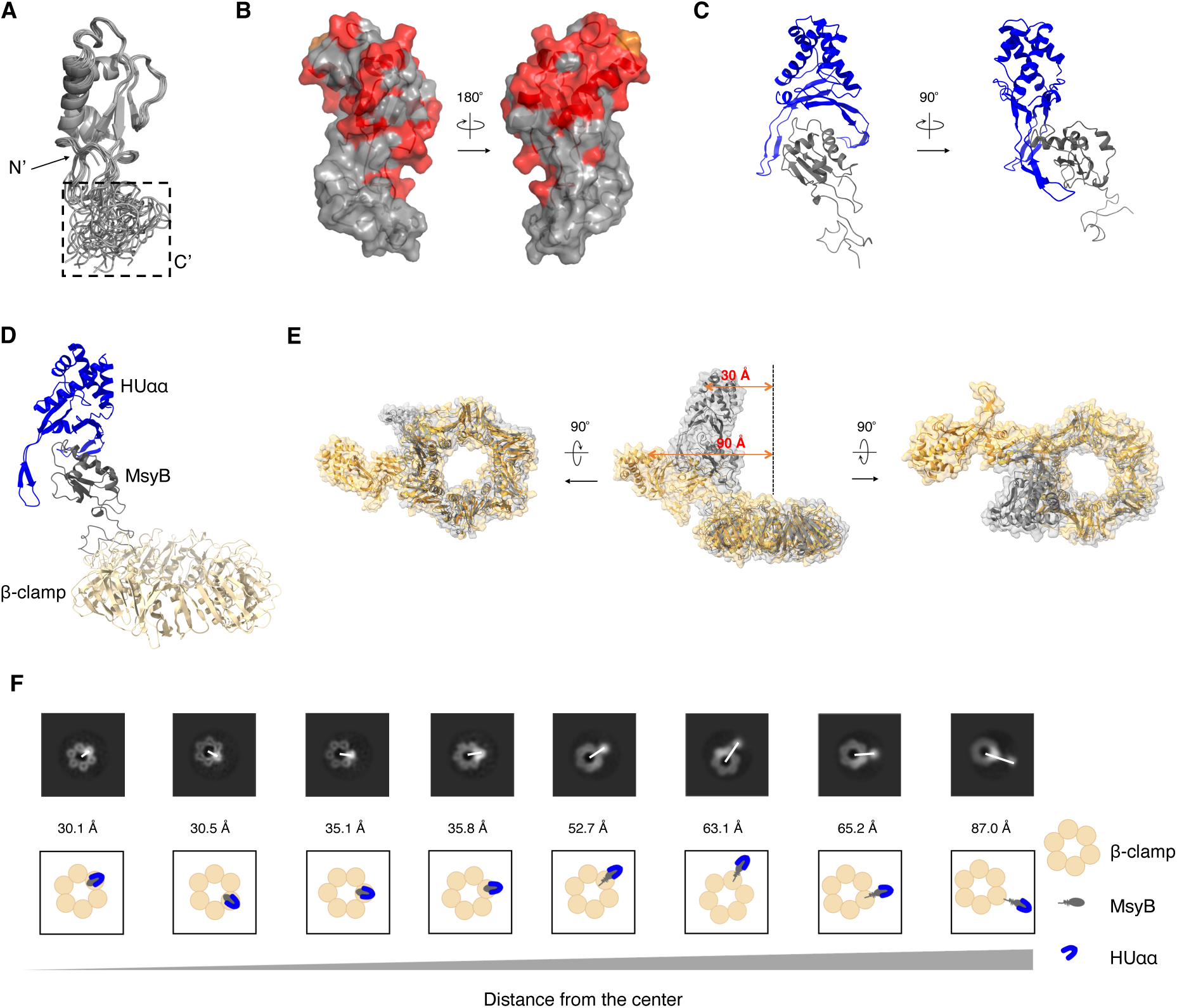
The HU-MsyB-clamp tripartite complex model. **(A)** The ensemble of 10 MsyB structures determined by NMR. The IDR domain is indicated by the dashed square. **(B)** The residues experiencing notable peak broadening effect (ratio < 0.2) as shown in fig. S9B are colored in red on the surface of MsyB with 180° rotation. P65 is colored in orange. **(C)** The cartoon illustration of HU-MsyB complex model built by HADDOCK. **(D)** The HU-MsyB-clamp tripartite complex model built by merging the HU-MsyB model and MsyB-clamp complex structure. **(E)** The overlay of the initial structure and final structure simulated for 10 ns in three different positions. **(F)** The 2D classification images of HU-MsyB-clamp complex showing the various distances between the outmost electron and the centre of the electron map. The cartoon illustration of the above images is also shown below.

To characterize the MsyB-clamp and HUαα-MsyB interactions, we performed NMR titrations utilizing the near-complete backbone assignment of MsyB (*62*) and tracked the residues which experienced peak broadening effect (Fig. S7C and D). In the MsyB-clamp titration spectra, MsyB residues Q38, V41, Q43, M48, H96, E97, D99, F103, L105 and E106 experienced the most notable peak-broadening effects with 1 molar equivalent β-clamp. Highlighting these residues on the structure of MsyB shows that they locate to the largest effect reside within the intrinsically disordered CTD (Fig. S7E). Q104 was not assigned and P107/P108 are not assignable albeit being in the same region (coloured in orange). A small region within the N-terminus (Q38-M48) is also affected which may suggest an allosteric link between the two. The contiguous region (F103-P108) within the C-terminal IDR form the major clamp-interacting surface, this region agrees with the cryo-EM structure, showing (F)QLEPP is the clamp binding sequence. Using MsyB^1-103^, a truncated mutant before the Q104, we repeated with the NMR titration. While MsyB^1-103^ is folded as the ^1^H-^15^N HSQC spectrum suggests, the titration showed no detectable interaction (Fig. S7F and G), showing that the N-terminal domain is not directly involved in the interaction. We then mutated the Q104 L105 to AA and a repeat NMR titration revealed no observable interaction with β-clamp (Fig. S7H). Complementarily, NMR titration shows no interaction between MsyB and mutant clamp^M362A, P363A^ representing mutations in the protein binding (Fig. S7I). Thus, the NMR and mutagenesis results agree with cryo-EM results, showing (F)QLEPP of MsyB and the protein-binding pocket of clamp is key for the interaction. Meanwhile, NMR titrations between MsyB and HUαα also revealed a clear interaction (Fig. S8A). The residues which experienced peak broadening effect in the titration was then mapped on the structure to locate the interface (Fig. 5B and S8B). The mapped structure suggests that NTD of MsyB is primarily involved in the interaction. The NMR complements the cryo-EM results with solution structure and the interface for the HUαα-MsyB interaction.

The cryo-EM and NMR structures, NMR titration between HUαα and MsyB, and the stoichiometry of the complex, allow us to build a HU-MsyB-clamp complex model. Firstly, using residues of MsyB that experienced most peak-broadening effect and DNA binding residues of HUαα as active residues, we built HUαα-MsyB model using HADDOCK (*63*). In the NMR-derived model, MsyB binds HUαα mainly on one side of saddle (Fig. 5C), similar to the position that of the known phage HU inhibitor Gp46 (*64*). Finally, the HUαα-MsyB model was merged with the EM structure to form a HUαα-MsyB-clamp complex tripartite model with the stoichiometry at 1:1:1 (Fig. 5D). As previous results suggest the existence of flexibility, the hybrid model was then subjected to molecular dynamics (MD) simulation for 10 ns to access the intrinsic elasticity and stability of the complex. During the simulation, the tripartite complex remained stable while MsyB swung away from the clamp ring, largely due to the ID domain of MsyB (Fig. 5E and Supplementary video 1).

Furthermore, the MD results also align with the 2D classification data, showing that HUαα swinging away from the clamp ring with maximum distance from the centre of HUαα to the centre of the ring on a 2D projection of 90 Å, similar to the measured value (87 Å) based on 2D classification (Fig. 5F). Therefore, these data suggest that the ID domain of MsyB is the main source of the elasticity observed within the tripartite complex.

### MsyB promotes adaptive mutation

As cells lacking HU are extremely sensitive to γ and UV irradiation (*65, 66*), removing the majority of HU from DNA by MsyB would significantly increase bacterial sensitivity to irradiation, and allowing active damage acquisition. As HU is mostly in MsyB-bound form during early stationary phase, MsyB could play a crucial role in bacterial stationary-phase adaptive mutation by removing large portion of HU from DNA. To verify this, we conducted a 30-day starvation experiment in which WT and Δ*msyB* strains were grown in LB medium for one month without additional nutrients (Fig. 6A). The final counts after 30 days differed significantly: 3.1 × 10^7^ CFU/mL for the WT and 3 × 10^3^ CFU/mL for the mutant (Fig. 6B). This result suggests a sginficantly reduced adaptability of the Δ*msyB* strain. Furthermore, whole-genome sequencing reveals that all four WT bacterial samples acquired two of the three shared single point mutations (SNP), whereas no mutations were found in the Δ*msyB* strain (Fig. 6C and SRR27561780 - 27561781 and SRR32051539 - 32051546). Among the three SNPs, one mutation was found in the cAMP-activated global transcriptional regulator (CRP), which is known to play a major role in catabolite repression during starvation (*67*). The results suggest that a sustained high level of MsyB allows bacteria to accumulate advantageous mutations, during which HU is removed from DNA by MsyB for damage acquisition. Therefore, the elevated MsyB population provides the first viable explanation for the previously unknown source of DNA damage in non-replicating cells.

**Figure 6.**
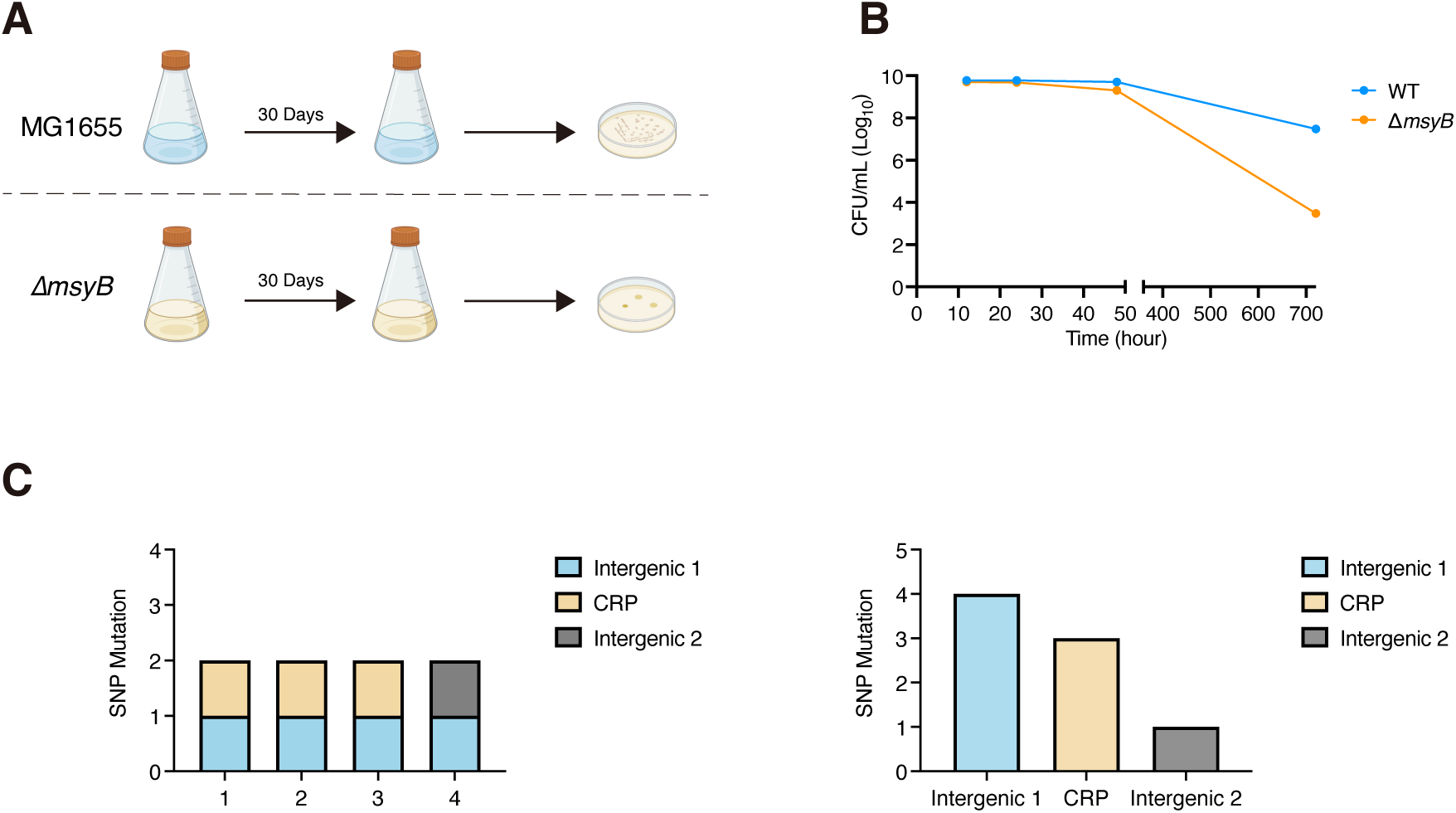
MsyB plays a significant role in *E. coli* stationary phase adaptive mutation. **(A)** The cartoon illustration of the 30-day starvation experiment. **(B)** The curves showing the CFU/mL of WT or Δ*msyB* determined at day 0, 1, 3 and 30. **(C)** Illustration of SNP mutations identified by whole-genome sequencing after the 30-day starvation experiment. Left: mutations identified in each colony; Right: distribution of the three different SNP mutations.

### Two-tiered HU-dependent stress responses that modulated by MsyB

Based on these results, we propose a mechanistic model for the two-tiered HU-dependent stress response modulated by MsyB (Fig. 7). Tier 1: When bacterial cells encounter a specific type of external stress, the cellular MsyB level is adjusted to release an appropriate amount of HU from the DNA. This adjustment likely coordinates with other stress-response pathways, altering DNA supercoiling thus gene expression to facilitate rapid stress adaptation. Tier 2: Under persistent stress that inhibits the growth of bacteria, such as long-term starvation, where transcript-level adaptation is insufficient, HU continues to be sequestered from DNA, allowing for the active acquisition of damage. Together with the β-clamp and Pol IV, mutations arise as a result of low-fidelity damage repair. Mutations that confer further adaptation are then selected by environmental pressure. Additionally, the MsyB-clamp interaction may facilitate the loading of the β-clamp onto exposed DNA, a prerequisite for Pol IV binding, but further investigation is required to support this hypothesis. Although adaptive mutation is not directly attributed to supercoiling, it represents an extension of bacterial stress responses linked to supercoiling regulation. To simplify the two responses within a unified MsyB-based model, we therefore group adaptive mutation under supercoiling regulation in this model. Notably, since MsyB is regulated by both σ^S^ and alarmone ppGpp (*68*), the HU-MsyB pathway is undoubtably overlapping with other pathways.

**Figure 7.**
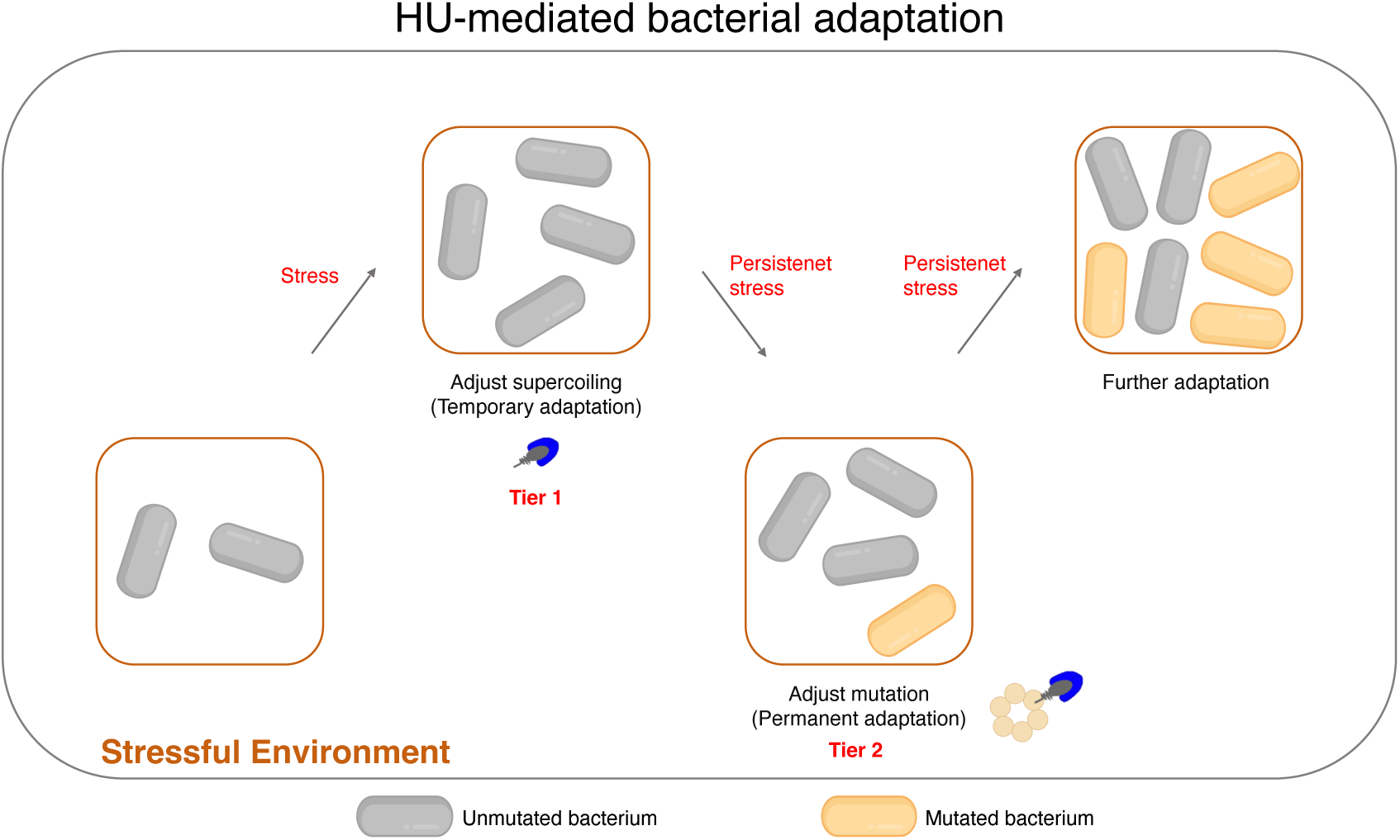
Two-tiered HU-dependent stress responses modulated by MsyB. A schematic illustration of the two types of stress responses regulated by MsyB through modulation of DNA-bound HU.

## Discussion

In this study, we report the HU-MsyB interaction as a novel regulatory mechanism for DNA supercoiling under stress conditions. This regulation is characterized by the fine-tuned expression of MsyB in response to various stresses. Using psoralen-based cross-linking and fluorescence imaging, we demonstrated changes in supercoiling profiles under different MsyB expression levels. Since psoralen preferentially interacts with unconstrained and unwound DNA while being blocked by nucleosomes (*51, 69*), this method does not detect all forms of supercoiling equally. However, it provides a sensitive approach to studying the HU-MsyB model, which is particularly responsive to psoralen-based cross-linking. Notably, under many stress conditions, DNA gyrase and topo I are often downregulated or their activities inhibited (*12–14*), suggesting that their role in stress response may be limited. While topoisomerases maintain supercoiling balance for essential DNA and RNA synthesis, our findings highlight the HU-MsyB interaction as a crucial mechanism in bacterial stress response.

During the stationary phase, the HU-MsyB interaction plays an additional role in inducing DNA damage within the compact nucleoid. The interaction between MsyB and β-clamp may also facilitate the recruitment of DNA Pol IV to DNA. Although HU composition shifts from HUαα to Huαβ during stationary phase, the affinity between HU and MsyB remains unchanged or may even be enhanced. This suggests that while HUαβ promotes nucleoid compaction during the stationary phase (*70*), the dynamic interplay between HUαβ-DNA and HUαβ-MsyB enables changes in nucleoid morphology as well as inducing DNA damage. Notably, IHF, another NAP that is known to constrain under- or over-twisted DNA (*53*), may also be removed from DNA by MsyB, particularly during the stationary phase when IHF and MsyB concentrations are high and HU levels are low.

The cryo-EM- and NMR-derived HU-MsyB-clamp complex model provides a mechanistic explanation for the two-tiered model: Strong HU-MsyB interactions promote the formation of the HU-MsyB complex over HU-DNA; While moderate MsyB-clamp interactions, flexibility of the complex and the vacant protein-binding pocket on the clamp ring likely facilitate subsequent proteins for clamp binding. As MsyB is highly conserved among all Gram-negative pathogenic bacteria (Fig. S9) and HU is crucial for bacterial responses to antibiotics, the HU-MsyB model is likely to apply to most Gram-negative bacteria. While the two-tiered HU-dependent stress response provides insights into supercoiling and adaptive mutations, its implications may be even more profound. Theoretically, since a specific level of supercoiling is required for an optimal stress response, misregulation of DNA-bound HU would leave bacteria in an improper physiological state for a given stress. Therefore, an exogenous HU inhibitor that removes the majority of HU from DNA would disrupt the finely tuned MsyB regulation necessary for precise stress adaptation, including adaptive responses for antibiotics and phage infection. Notably, Bacillus phage SPO1 produces the HU inhibitor Gp46 to optimize infection of its host (*64, 71*). While bacterial topoisomerases have long been among the most important antibiotic targets and continue to attract considerable attention, HU, with its other roles in cell viability and biofilm formation, is therefore another potential drug target for antibiotic development.

## Supporting information

move s1

## Acknowledgement

The authors would like to thank the NMR centre of the first affiliated hospital of Xi’an Jiaotong University for the spectrometer time, and centre for instrumental analysis and metrology of Wuhan institute of virology for providing technical assistance and cryo-EM data acquisition. Ms Zonglan Yu provided technical support for the gene knockout assays.

## Data availability

The cryo-EM maps associated with this study have been deposited in the Electron Microscopy Data Bank under accession codes EMD-61841. Structure coordinates have been deposited in the RCSB Protein Data Bank with accession codes 9JVO. The sequencing data for the WT and Δ*msyB* strains in the adaptative mutation experiment is available under SRR27561780 - 27561781 and SRR32051539 - 32051546. The RNA-seq data are available in the SRA under accession number: PRJNA1227296.

## Author Contributions

B.L. designed the experiments. B.L., S.Y., and H.W. conceived the project. H.C, Y.Y, X.C, H.Z, Y.H., M.L., W.L., Z.Y., and Y.W. performed the experiments. B.L. analysed the data. B.L. wrote the first draft of the manuscript with contributions from all authors.

