## Supplemental files for "MsyB-HU Interaction Modulates a Two-Tiered Bacterial Stress Response by Regulating DNA Supercoiling"

### Material and methods

*Recombinant protein expression*

All plasmids were constructed using Gibson-assembly. The *hupA*, *hupB*, *ihfA* and *ihfB* genes were chemically synthesized while other target genes were amplified directly from bacterial genomes by PCR. The PCR products were then purified by 1.5% agarose gel electrophoresis and cloned into relevant linearized vectors. The *ihfA* and *ihfB* genes were cloned into pETDuet-1 vector and *dnaN* gene was cloned into pET-46 vector, whilst the rest were cloned into pET-SUMO vector. The *ihfA* contained an N-terminal $His_6$ tag, followed by a TEV protease cleavage sequence. Site-directed mutagenesis was performed by back-to-back PCR amplification using Q5 Site Directed Mutagenesis Kit (NEB, Cat#E0552S). All primers and plasmids used in the study are listed in Table S1. All constructs were confirmed by DNA sequencing. The relevant vectors were transformed into *E. coli* BL21(DE3) strain for the expression of the recombinant proteins. Bacteria culture were incubated at 37°C in LB medium or M9 medium ($^{15}N$-labelled $NH_4Cl$ and/or $^{13}C$-labelled glucose were used for isotopic labelling) containing relevant antibiotics till $OD_{600}$ of ~ 0.6 and induced by 1 mM IPTG. The cell cultures were then harvested by centrifugation at 5,000 rpm for 10 min at 4°C after incubation at 18°C for 16-18 hours and resuspended in a 30 mL binding buffer (50 mM $NaH_2PO_4$, 300 mM NaCl, 10 mM imidazole and pH 8.0). The harvested bacterial cultures were lysed by sonication. The lysate was then centrifuged at 20,000 rpm for 30 min at 4°C and the supernatant fraction was subjected to immobilized metal affinity chromatography using the Ni-NTA agarose resin for standard his-tagged protein purification (Qiagen, Cat#30250). The resin was washed in a wash buffer (the binding buffer supplemented with 20 mM imidazole), and the recombinant proteins were eluted in a 30 mL elution buffer (the binding buffer supplemented with 250 mM imidazole). The purified proteins were dialyzed into a storage buffer (50 mM NaCl, 20 mM Tris, 0.2 mM TCEP and pH 7.4) at 4°C overnight. TEV protease was added to IHF to remove the $His_6$ tag, whilst the SUMO tag was removed by adding SUMO protease Ulp1. The flow-through was passed through Ni-NTA resin twice to separate the residual protease and the fusion tag. To get DNA-free NAPs, proteins were loaded on a 5 mL HiTrap Heparin HP column (GE Healthcare Life Sciences), eluting with a linear gradient of 0.05-1.5 M NaCl. The recombinant proteins were further purified by size exclusion chromatography using a Superdex 200 Hiload 16/600 column (GE Healthcare Life Sciences), which were equilibrated with the storage buffer, on an ÄKTA pure system (GE Healthcare Life Sciences). The concentrations of HU and IHF proteins were determined by the Bradford protein assay, while the concentrations of the other proteins were determined by measuring absorbance at 280 nm using a nanodrop. HUαβ heterodimers were prepared by mixing equal molar amount of HUαα and HUββ homodimers.

*BLI analysis*

Protein-protein binding kinetics were measured by a double-channel Octet RED (Sartorius) using Ni-NTA biosensors (Sartorius, Cat#18-5101) at 30°C with shaking at 1,000 rpm. The sensors were pre-hydrated for 10 min in the BLI buffer (the storage buffer containing 0.02% Tween 20). The N-terminal $His_6$-tagged β-clamp (100 nM) was immobilized on the probes for 180 s, followed by a 60 s baseline measurement. The sensors were immersed in MsyB for 5 min with the concentrations ranging from 4.2 μM to 66 μM for association. For the affinity between HUαα-MsyB and β-clamp, the sensors were dipped in the wells containing the HUαα-MsyB complex at various concentrations (1.4-21.7 μM). Dissociation was performed in the same BLI buffer for 10 min. The $K_D$ was determined by fitting the reference-subtracted sensorgrams globally with the 1:1 binding model using the Octet analysis software provided by the manufacturer. For the MsyB-NAPs interaction, 125 nM of C-terminal $His_6$-tagged MsyB was used and the rest steps followed the protocol described above.

*BACTH assay*

A bacterial two hybrid system based on adenylate cyclase reconstitution (Euromedex, France) was used to study protein-protein interactions *in vivo.* The *msyB* gene were cloned into pKT25 or pKNT25 plamsid. Genes encoding β-clamp and HUα were respectively cloned into pUT18 or pUT18C plasmid. These plasmids encode hybrid proteins with the T18 or T25 domains of adenylate cyclase fused to either the N-terminal or C-terminal of the target protein. All constructs were confirmed by DNA sequencing. To test protein-protein interactions, a pKT25 or pKNT25 derivative and a pUT18 or pUT18C derivative were co-transformed into the adenylate cyclase-deficient *E. coli* strain BTH101 competent cells by electroporation. The transformed cells were then plated onto LB plates containing 100 μg/mL ampicillin, 50 μg/mL kanamycin, 40 μg/mL X-gal (5-bromo-4-chloro-3-indolyl-d-galactopyranoside) and 0.5 mM IPTG. pKT25-zip and pUT18C-zip co-transformants were used as the positive control, while pKT25 and pUT18 co-transformants were used as the negative control. Bacteria were grown at 30°C for 36 hours and blue colonies were then quantified by measruring the β-galaxtosidase activity using o-nitrophenol-d-galactopyranoside (ONPG) as a substrate. For this assay, transformants were cultured in LB broth supplemented with antibiotics and induced with 1 mM IPTG. After incubation at 30°C for 36 hours, the cells were lysed with chloroform and SDS and the β-galactosidase activity was determined.

*Pull-down assay and MS analysis*

Pull-down assays were performed using Ni-NTA resin as describe previously (*1*). Briefly, 4 litres of *E. coli* MG1655 culture were harvested at $OD_{600}$ of ~ 0.6 by centrifugation at 5,000 rpm at 4°C for 10 min, resuspended in 50 mL of the binding buffer (50 mM NaCl, pH 8.0, 25 mM $NaH_2PO_4$, 10 mM imidazole and 5% glycerol) and lysed by ultrasonication. After being centrifuged at 20,000 rpm for 30 min at 4°C, the supernatant of the lysate was loaded onto the Ni-NTA resin with C-terminal $His_6$-tagged MsyB immobilized. The column was then incubated at 4°C for 2 hours before proceeding to the standard protein purification protocol. Pull-down elution sample was first analyzed by gel electrophoresis and sample bands recycled from the SDS gel were digested with the trypsin and desalted by C18 ZipTips (Millipore, Cat# ZTC18S096). The samples were analyzed by a Q-Exactive plus mass spectrometer (Thermo Fisher Scientific) coupled with an UltiMate 3000 RSLCnano system. Database search was performed by MaxQuant (*2*).

*EMSA*

The EMSA assays were performed with dsDNA (the sequence is listed in Table S1), MsyB, and HUαα or HUαβ in the storage buffer containing 1 mM EDTA and 5% glycerol. HUαα or HUαβ was incubated with dsDNA on ice for 30 min as illustrated in Fig. 1F and G. After adding MsyB, the mixtures were incubated on ice for another 10 min before loading onto 0.75% agarose gels in 0.5 × TBE. The electrophoresis was performed at 90 V in 0.5 × TBE buffer for 90 min on ice. Images were captured by a ChemiDoc$^{TM}$ MP Imaging System (Bio-Rad).

*RT-qPCR*

The WT strain was grown in LB medium to mid-logarithm phase. Then, the cells were used to inoculate (1% v/v) into fresh LB medium, LB with 1 μg/mL of MMC, M9 medium, M9 with 1 μg/mL of MMC, M9 medium (pH 6, adjusted with HCl), LB medium with antibiotics (10 μg/mL of CB, 1 μg/mL of TET, or 0.2 μg/mL of LFX) respectively. 1 μg/mL of MMC was added to the LB culture at 13 hours. The cultures continued to be incubated at 37°C or 42°C for 24 hours, and samples were took out as indicated.

Total RNA was extracted from each sample using the RNeasy® Mini Kit (QIAGEN, Cat#74104), according to the manufacturer's instructions. 1 μg of total RNA was reverse transcribed with qPCR RT Master Mix Kit (TOYOBO, Cat#FSQ-301) for complementary DNA synthesis. RT-qPCR was performed in a 20 μL volume consisting of 10 μL of 2 × QuantiNova SYBR Green PCR Master Kit (QIAGEN, Cat#208054), 6.2 μL of RNA-free water, 1 μL of cDNA template (containing 0.5 μg cDNA), and 1.4 μL of each primer pair according to the manufacturer's instructions using the CFX Connect Real-Time PCR System (Bio-Rad). All experiments were carried out three times. The specific primers used for each gene are listed in table S1. Relative differential expression levels of *msyB* were calculated based on the cycle threshold ($C_T$) values normalized with the reference gene (16S ribosomal RNA gene) using the $2^{-\Delta\Delta CT}$ method.

MsyB$^{OE}$, *and ΔmsyB and ΔhupAB strains construction*

The MG1655 bearing the pBAD-MsyB plasmid was induced with 0.2% L-arabinose to overexpress MsyB. The *ΔmsyB* strain was generated by the CRISPR-Cas9 system as previously reported (*3*). Briefly, the pTargetF (miaolingbio, Cat#P0743) was linearised by inverse PCR. A designed N20 sequence of *msyB* with overhangs was amplified by PCR and ligated into linearised pTargetF to obtain pTargetF-*msyB*. The donor DNA fragment was formed by overlap extension PCR, in which two fragments flanking *msyB* were combined. 100 ng of pTargetF-*msyB* and 400 ng of donor DNA fragment were co-electroporated into competent *E. coli* MG1655 carrying the pCas plasmid (miaolingbio, Cat#P0762), which expresses Cas9 protein and the λ-Red recombination system. Cells were selected on LB agar medium containing kanamycin and spectinomycin at 30°C overnight, and verified by colony PCR and DNA sequencing. The correct mutants were treated with 0.5 mM IPTG for 8-16 hours to cure the pTargetF-*msyB* and cultivated at 37°C overnight to cure the temperature-sensitive pCas. The *ΔhupA* strain was obtained using the same protocol, followed by the construction of the *ΔhupAB* strain based on the *ΔhupA* strain. Primers used for these constructs are listed in Table S1.

*Bacterial growth curves determination*

The WT and *ΔmsyB* strain were grown in LB medium overnight before diluted 100 times into fresh medium like LB medium, M9 medium, LB with 1 μg/mL of MMC, M9 with 1 μg/mL of MMC, M9 medium (pH 6, adjusted with HCl), LB medium with 10 μg/mL of CB, LB medium with 5 μg/mL of CB, LB medium with 1 μg/mL of TET, or LB medium with 0.2 μg/mL of LFX. The bacterial growth assays were performed at 37°C or 42°C in 96-well plates. The absorbance at 600 nm was measured every hour using a Neo 2 multi-well plate reader (BioTek). At least three biological and technical replicates were performed for each growth curve test.

*bTMP incorporation and imaging*

Cells were harvested at $OD_{600}$ of 0.4-0.5 and 1.0-1.2 for exponential phase and stationary phase experiment, respectively. Bacterial cells were treated as indicated and washed with PBS before they were permeabilized with 250 μg/mL lysozyme for 30 min at room temperature (RT). Cells were incubated with 0.5 mg/mL EZ-Link Psoralen-PEG3-Biotin (ThermoFisher Scientific, Cat#29986) for 20 min, then exposed to 4 kJ $m^{-2}$ of 365 nm light using an UV Crosslinker for 20 min on ice followed by a PBS wash. Cells were then fixed with 4% paraformaldehyde and treated with 0.2% Triton X-100 before they were incubated with Alexa Fluor 594 Streptavidin (ThermoFisher Scientific, Cat#S11227) for 1 hour at RT in the dark, washed with PBS and stained with DAPI for 10 min. Cells were then mounted onto glass slides with ProLong Gold antifade mountant (ThermoFisher Scientific, Cat#P36934). Images were taken with Olympus FV3000 confocal microscope (Olympus). Fluorescence intensity was quantified using ImageJ software from multiple random fields obtained with identical acquisition settings and normalized by cell number.

*Gram staining and Methylene blue staining*

Bacteria were harvested by centrifugation at 4,000 rpm for 10 min and washed twice with PBS buffer. The cells were resuspended in PBS and 5 μL of the bacterial suspension was spread onto clean glass slides to create thin films. The slides were passed through a flame 3-4 times to heat-fix the bacteria to the slide. For Gram staining, the slides were strained firstly with crystal violet solution for 1 min followed by Gram's iodine solution for another 1 min. 95% ethanol was used to decolorize the slides and the safranin solution was used as counterstain. For methylene blue staining, samples were strained with a 0.1% methylene blue solution for 1 min and then gently rinsed with distilled water. The slides were air-dried and covered with cover slips. Images were taken with a CX23 upright microscope (Olympus).

*Transcription profiling*

For RNA-Seq, WT and Δ*msyB* strains were first grown overnight in LB medium and then overnight cultures were diluted 1:100 in fresh regular LB medium or LB with 10 μg/mL of CB. Cells were harvested at $OD_{600}$ of ~ 0.5-0.6 (exponential pahse) or 1.0-1.2 (stationary phase). Three biological replicates per sample were collected for each condition. Total RNA was extracted using the RNeasy Mini Kit (Qiagen) and ribosomal RNA was removed with Ribo-Zero Plus rRNA Depletion Kit (Illumina, Cat#20040529). The remaining RNA was fragmented, converted to cDNA (with dUTP incorporation for strand specificity), and used to create a sequencing library. Library quality was assessed with qBit, Agilent Bioanalyzer and qPCR. Sequencing was performed on a PE 150 system (Illumina). Reads were mapped to the *E. coli* MG1655 reference genome using Bowtie2, and gene expression was quantified using FeatureCounts (normalized to RPKM). Differential gene expression was determined with DESeq2 (fold change > 1, adjusted $p < 0.05$). KEGG pathway enrichment analysis was performed using clusterProfiler. The RNA-seq data are available in the SRA under accession number: PRJNA1227296.

*cryo-EM sample preparation and data collection*

HUαα-MsyB complex was prepared firstly. Excess MsyB was mixed with HUαα in the storage buffer at 4°C overnight. Redundant MsyB was separated from HUαα-MsyB complex using a Superdex 200 Hiload 16/600 column. Then $His_6$-tagged β-clamp was added to the HUαα-MsyB complex and the mixture was dialyzed into cryo-EM buffer (20 mM Tris, 0.1 mM $EDTA·Na_2$, 0.2 mM TCEP and pH 7.4). HUαα-MsyB-clamp complex was purified by size exclusion chromatography again.

For cryo-EM sample preparation of the HUαα-MsyB-clamp complex, we applied 4 μL of the mixed proteins was applied to glow-discharged gold (Au) 200 mesh R1.2/1.3 holey carbon grids (Quantifoil Micro Tools GmbH). The grids were incubated for 20 s, followed by blotting for 2 s using a blot force of 5 in an environment maintained at 100% humidity and 4°C. Subsequently, the samples were plunge-frozen using Vitrobot Mark IV (ThermoFisher Scientific).

Data collection was performed using a CRYO ARM 300 electron microscope (JEOL) operated at an acceleration voltage of 300 kV. Imaging was done with a K3 direct electron detector (Gatan) at a nominal magnification of 50,000 × in super-resolution counting mode. This setup yielded a super-resolution pixel size of 0.475 Å/pixel. Movies were collected automatically via Serial-EM software (*4*) at 40 frames per second, achieving a total electron dose of 40 e/Å². Data were collected within a defocus range of -0.5 to -2.5 μm.

*Cryo-EM data processing*

For the HUαα-MsyB-clamp complex dataset, recorded videos were processed using cryoSPARC v.4.4.1 (*5*). We began with patch motion correction and then performed contrast transfer function (CTF) estimation. Poor-quality images were identified and discarded. From 14,140 micrographs, a total of 3,621,808 particles were automatically picked and underwent two rounds of 2D classification to remove unsuitable particles. The 804, 868 particles that exhibited clear class averages were used for ab initio reconstruction and heterogeneous refinement. Following this, CTF refinement was applied to the most prominent class. We then performed non-uniform refinement to optimize the density map, and local resolution estimation was carried out to assess the quality of the reconstructed image.

*Model building and refinement*

To construct the structure model of the complex, initial model β-clamp (PDB:1OK7) was derived from the Protein Data Bank (PDB) (*6*). The initial model was rigid body fitted into the cryo-EM map in UCSF ChimeraX (*7*). Further adjustments and MsyB peptides model building were manually made in Coot (*8*). Comprehensive real-space refinements were carried out using Phenix (*9, 10*). Details on data validation and statistical outcomes are provided in the supplementary materials.

*NMR Structure Determination and titration*

The NOE experiments were recorded on a 900 MHz Avance III spectrometer (Bruker) at 298 K equipped with a cryo-probe. Two 3D $^{1}H$-$^{15}N$ and $^{1}H$-$^{13}C$ NOESY-HSQC experiments and the chemical shift assignment (BMRB ID: 28093) were used in the final structure calculation. The ARIA (v2.3) was the software for automated NOE assignment and structural calculation. The tolerances for NOE assignment were ± 0.05 ppm for both direct and indirect proton dimensions, ± 0.5 ppm for nitrogen and ± 1 ppm carbon dimensions, respectively. The ARIA parameters p, Tv, and Nv were set to default values. One hundred ninety-seven dihedral angle restraints derived from TALOS+ were also implemented. The final 10 lowest energy structures had no NOE violations greater than 0.5 Å and dihedral angle violation. The data were processed by NMRpipe (*11*) and analyzed using NMRView (*12*). The titration experiments were performed in the storage buffer. A series of $^{1}H$-$^{15}N$ HSQC spectra were recorded at 298 K using an 800 MHz Avance Neo spectrometer (Bruker) equipped with a cryogenic probe.

*Molecular Docking using HADDOCK*

Using amino acids of MsyB that experienced most peak-broadening effect and those of HUαα revealed by NMR titration experiment, the NMR structure of MsyB and structure of HUαα (homology modelling based on 8HD5) were used to build a model of a 1:1 complex of HUαα-MsyB-clamp complex using *HADDOCK* (v2.4) (*13*). Amino acids experienced more than 50% intensity reducation were assinged as active residues, and the surrounding residues were designated as passive residues. An ambiguous distance restraint of 2.0 Å was invoked between all active and passive residues of the other protein partner. The interfacial residues were allowed to move during the simulated annealing and water refinement. A total of 1,000 initial complex structures were generated by rigid-body energy minimization, and the best 200 by total energy was selected for torsion angle dynamics and subsequent Cartesian dynamics in an explicit water solvent. The best structure was chosen based on interactional energy.

*MD simulation*

MD simulations were performed for meticulously assessing the flexibility and stability of the HUαα-MsyB-clamp complex, ensuring a comprehensive representation of the biological system. The initial model of the HUαα-MsyB-clamp complex was built by docking HUαα and MsyB to the β-clamp ring according to the position of the peptide of MsyB in the cryo-EM structure we got. The MD simulations were set up using CHARMM-GUI (*14*) solution builder, allowing for the comprehensive generation of the MD input. Using the TIP3P model to represent water molecules. To maintain electrostatic neutrality within the complexes, 150 mM NaCl was added employing the Monte Carlo Method. The force fields employed were CHARMM36m for the system, while CHARMM General Force Field (CGenFF) generated the ligand force field parameters (*15*). The systems underwent an initial energy minimization phase comprising 20,000 steps utilizing Gromacs version 2023.3 (*16*), followed by a controlled heating process, gradually raising the temperature from 0 K to 298.15 K in the NVT ensemble over a simulation period exceeding 1000 ps, maintaining conservation of particles, volume, and temperature. Further simulations were conducted under a constant pressure of 1 atm in the NPT ensemble, conserving particle numbers, pressure, and temperature, employing 10.0 kcal $mol^{-1}$ $Å^{-2}$ harmonic restraints. The simulations, including both the NVT and NPT ensembles and the subsequent MD production employing a 2fs time step for 10 ns. The comprehensive analysis of simulation outputs and structural features was carried out utilizing Gromacs (*16*), ChimeraX (*7*), Pymol (*17*) and VMD (*18*).

*Short-term stationary phase experiments and sequencing*

Three populations of WT or Δ*msyB* strain were initiated from single colonies. The bacteria were inoculated into six 2.5-liter polycarbonate breathing flasks containing 1 L LB media in an incubator at 37˚C, while shaking at 220 rpm for 30 days. No new nutrients or resources were added to the cultures with time, except for sterile water that is added once to compensate for evaporation at 15 days, according to the weight lost by each flask during that time period. 1 mL of the cultures were taken out and were serial diluted before plated on LB agar plate and grown overnight at 37°C to determine the cell counts (CFU/mL). The starting single colonies, and six colonies from WT or Δ*msyB* group that harvested at day 30, were prepared for whole genome sequencing. Whole genome sequencing was performed on an HiSeq2000 platform (Illumina). And the data was analyzed using the method adapted from previous reports (*19, 20*). Trimmomatic software (*21*) was used to filter out low-quality reads from the raw data. Bowtie2 software (*22*) was employed to map the filtered data to the bacterial genome at hour 0. Variants were identified from uniquely mapped reads using the mpileup function in SAMtools (-Q 30) (*23*). When calculating single-point mutations and total matched base pairs, bases with a coverage depth <50 × were discarded. Additionally, only mutations with a mutation rate > 80% at a single point were considered valid. The overall mutation rate was calculated by dividing the sum of variant base coverage at all valid mutation sites by the sum of base coverage at all valid base coverage sites on the genome.

*Phylogenetic analysis*

To perform the phylogenetic analysis, the msyb gene sequence was retrieved from the NCBI database, and the top 5000 sequences were downloaded. These sequences were then aligned using Mafft v7.515 with the "-auto" parameter. The resulting alignment was used to construct a phylogenetic tree using FastTree v2.1.11 and visualized with Chiplot (*24*).

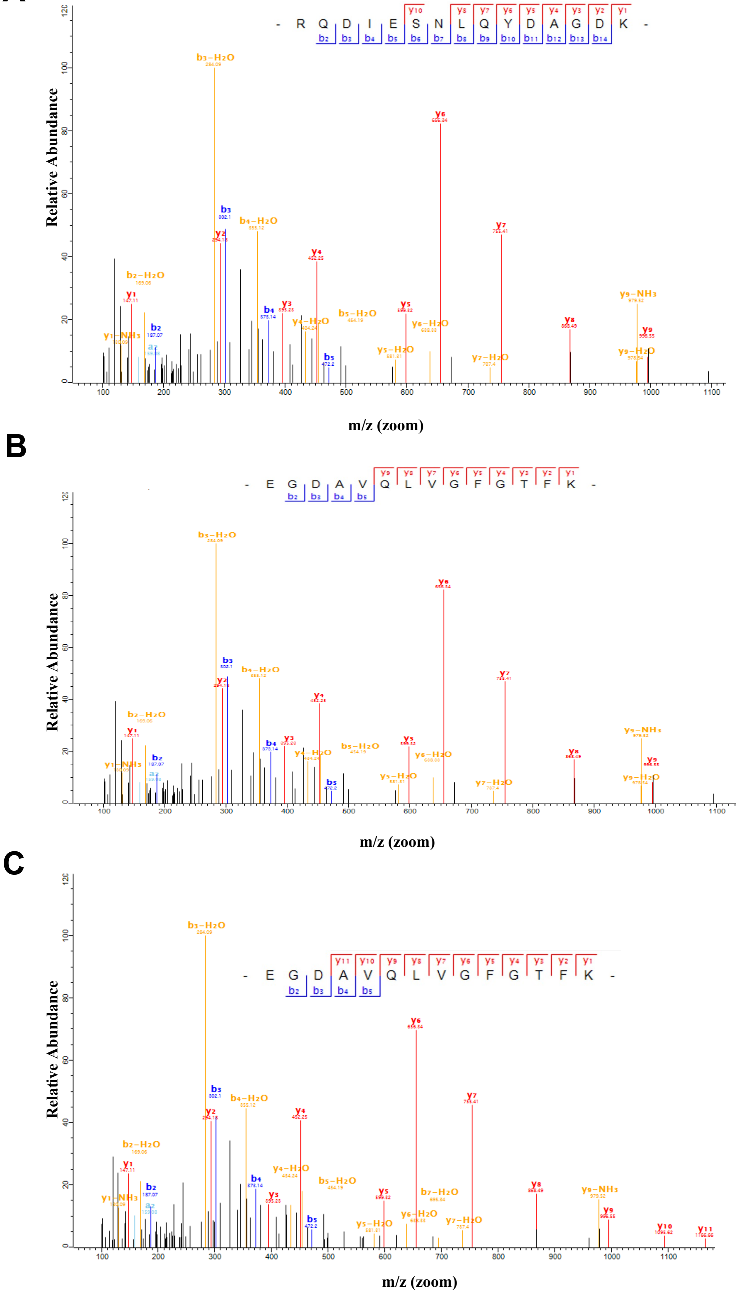

**Supplementary figure 1: Protein identification by Mass spectroscopy**
(**A-C**) Representative MS/MS spectra of unique peptide (RQDIESNLQYDAGDK ) for GlmS (the 65 kDa band ), EGDAVQLVGFGTFK for HUα (the 22 kDa band) and EGDAVQLVGFGTFK for HUα (the 10 kDa band) identified from trypsin digested pull-down proteins by a Q-Exactive mass spectrometer, respectively.

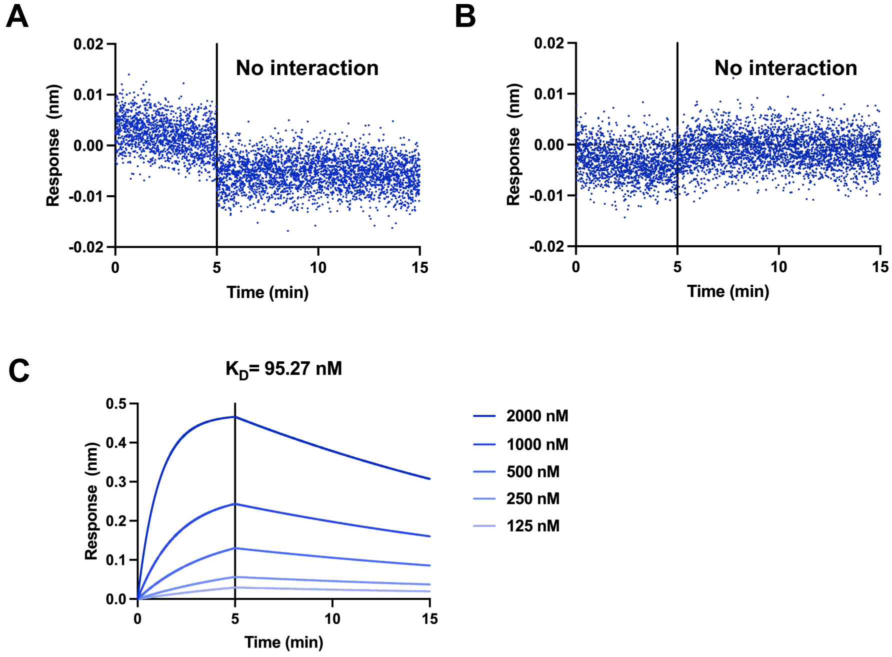

**Supplementary figure 2: The interactions of MsyB between other major NAPs.** (**A-C**) The affinity determination by BLI for MsyB-FIS, MsyB-H-NS and MsyB-IHF interactions.

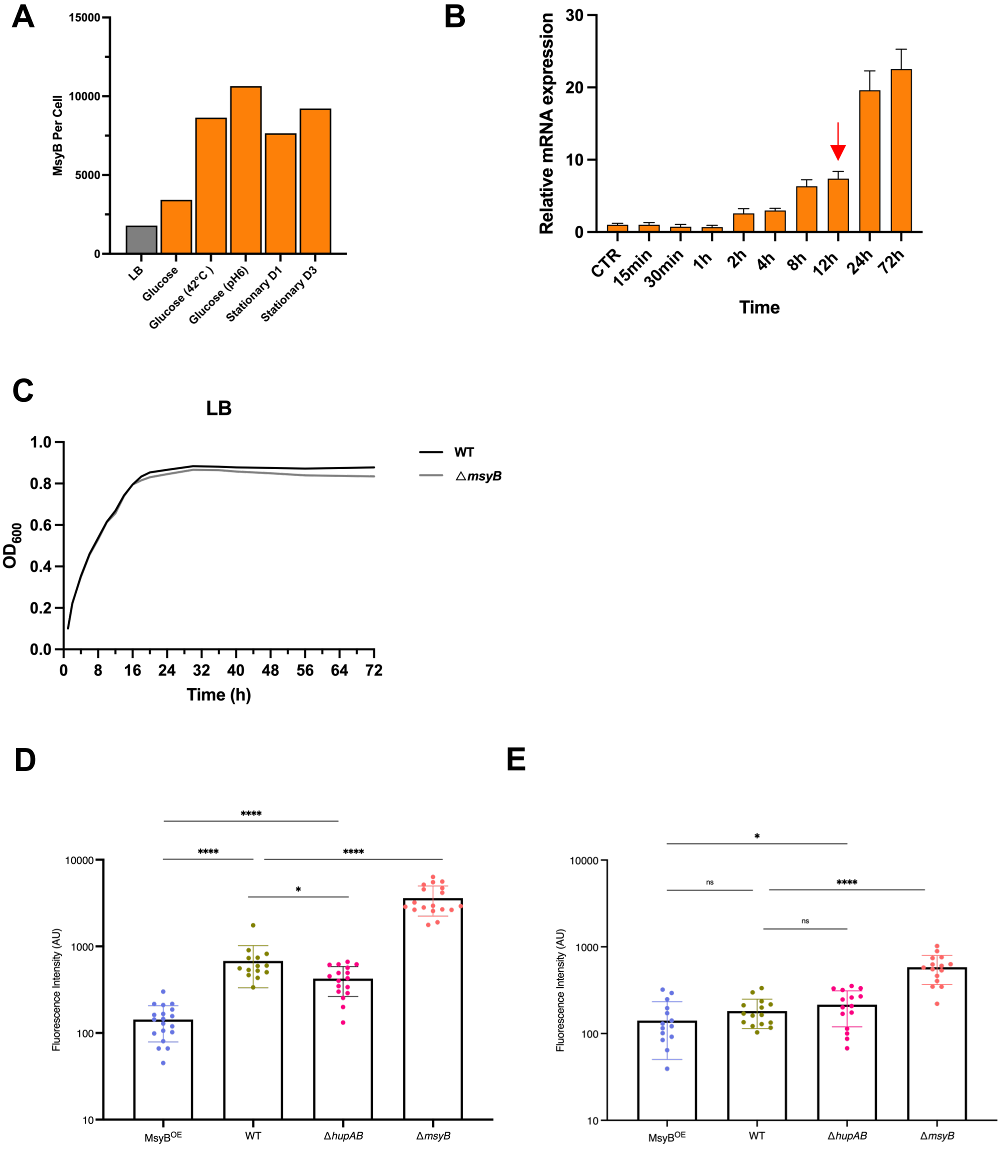

**Supplementary figure 3: MsyB inhibits DNA supercoiling.**
(**A**) The population count of MsyB (number per cell) under different stress conditions.
(**B**) The relative expression levels of *msyB* at different time points, from the exponential to stationary phase, were determined by q-RT-PCR. The arrow indicates the start of the stationary phase after approximately hour 12. (**C**) The 72-hour growth curves of WT and Δ*msyB* strains from the growth phase to the stationary phase. (**D-E**) Quantification of supercoiling for MsyB$^{OE}$, WT, Δ*hupAB* and Δ*msyB* cells during the exponential and stationary phases. Each point represents the integrated fluorescence signal from a randomly chosen field. Values are expressed as mean ± SD and normalized to cell number. *$P < 0.05$ and ****$P < 0.0001$. For qPCR and growth analysis, three biological replicates were performed, each containing three technical replicates. The growth curve represents the average $OD_{600}$ from a representative experiment.

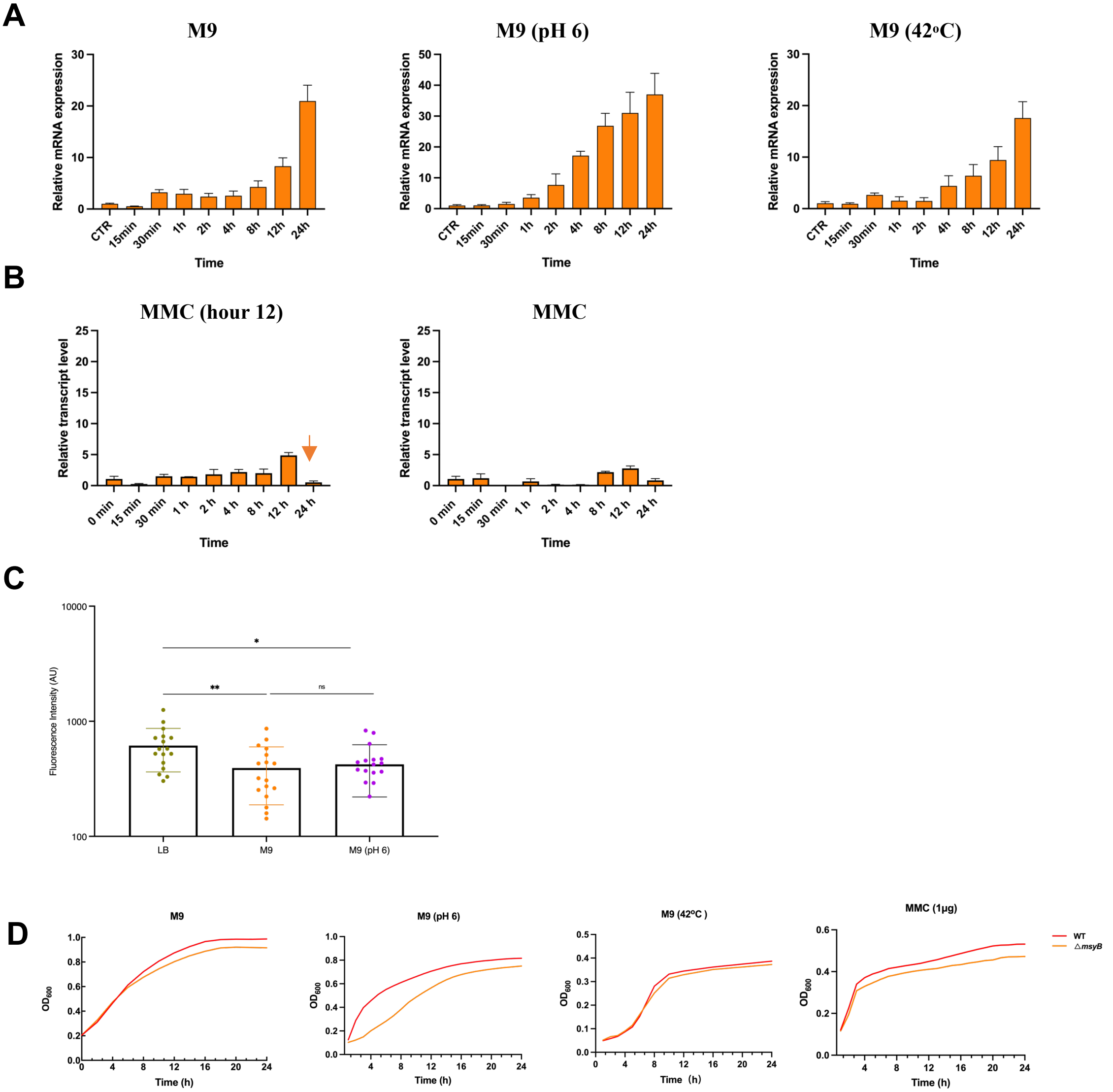

**Supplementary figure 4: MsyB modulates DNA supercoiling under various stressful conditions**

(**A**) The *msyB* transcript level determined by q-RT-PCR at time points shown for three different stresses: nutrient limitation, acidic pH and 42°C. (**B**) The *msyB* transcript level determined by q-RT-PCR during MMC-induced SOS response. Left: .MMC was added to WT at the start of the stationary phase. Arrow points the start of stationary phase when MMC is added. Right: MMC was added in the beginning of the incubation. (**C**) The quantification of supercoiling for WT cell at nutrient limitation (M9: glucose as the only carbon source) and acidic condition. Each point denotes the integrated fluorescence signal from a randomly chosen field. Values are mean±SD and are normalized with cell number. . *P < 0.05 and **P < 0.01 *** *P*<0.001. (**D**) The growth curves of the WT and *ΔmsyB* strains showing the impaired growth in the *ΔmsyB* strain under M9, M9 (pH 6), M9 (42°C) and MMC-treated conditions. For qPCR and growth analysis, three biological replicates were performed, each containing three technical replicates. The growth curve represents the average $OD_{600}$ from a representative experiment.

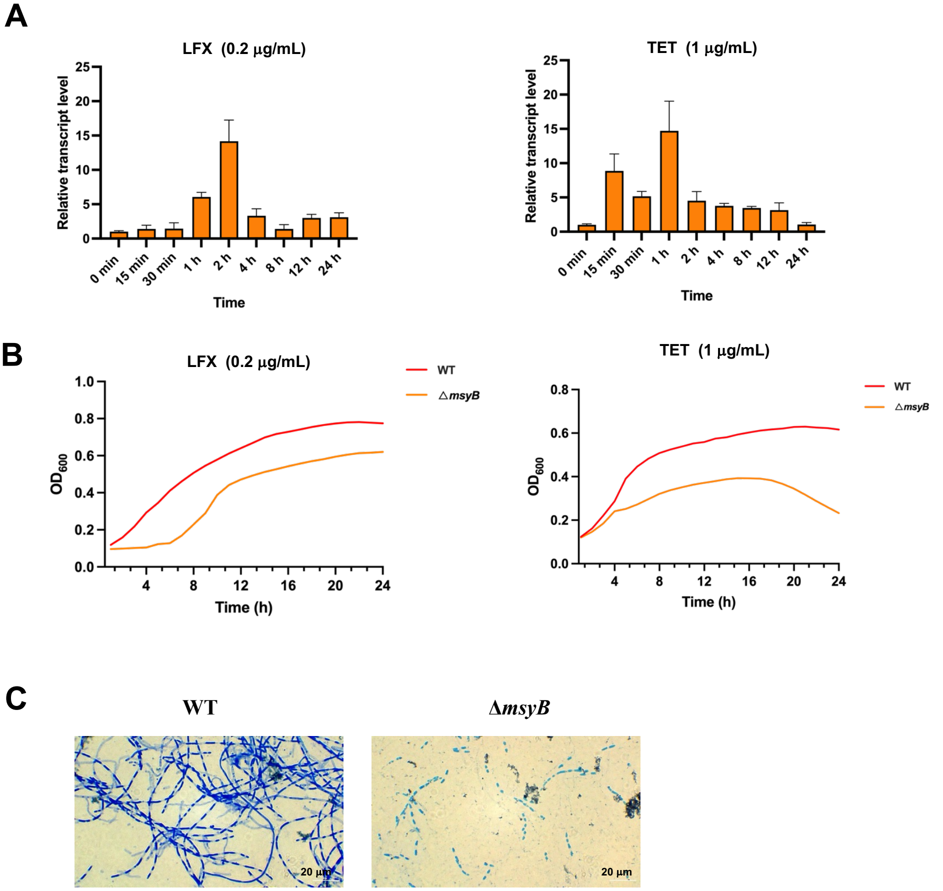

**Supplementary figure 5: MsyB modulates DNA supercoiling under antibiotic treatment** (**A**) The *msyB* transcript level determined by q-RT-PCR at time points shown under LFX (Left) and TET (right) treatment. (**B**) The growth curves of the WT and *ΔmsyB* strains showing the impaired growth in the *ΔmsyB* strain under LFX and TET treatment. (**C**) Methylene blue staining of the samples in Fig. 2G, showing the most of the cells in the *ΔmsyB* group were dead. Scale bar: 20 **μ**m.

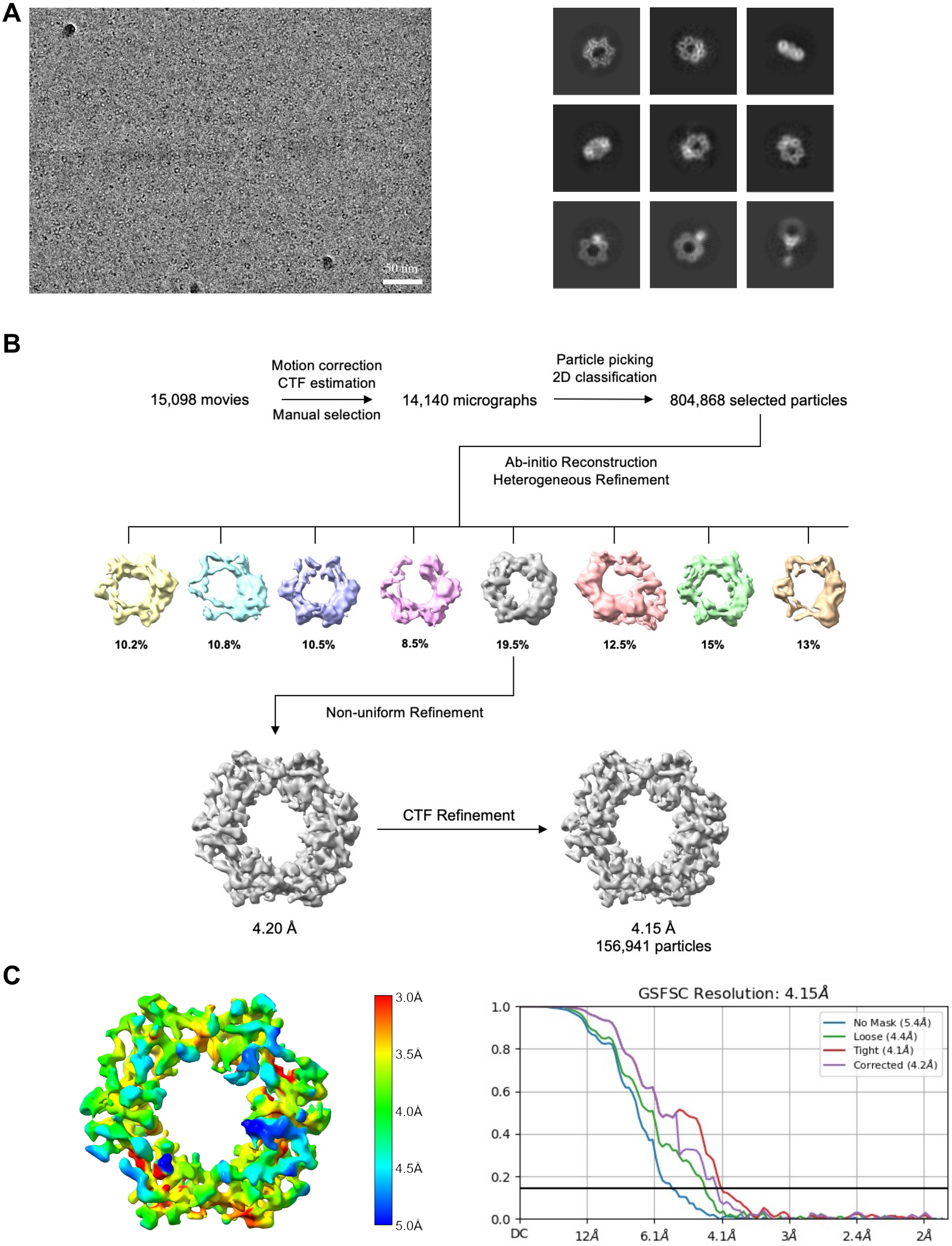

**Supplementary figure 6: Cryo-EM analysis of the clamp-MsyB-HUαα Dataset**
**(A)** A representative motion-corrected micrograph and reference-free 2D class averages generated using cryoSPARC. **(B)** A schematic of the workflow used for 3D reconstruction.
**(C)** Local resolution distribution and the plots of the Gold Standard Fourier Shell Correlation (GSFSC) of the final map used for model building from CryoSPARC with a reported resolution of 4.15Å.

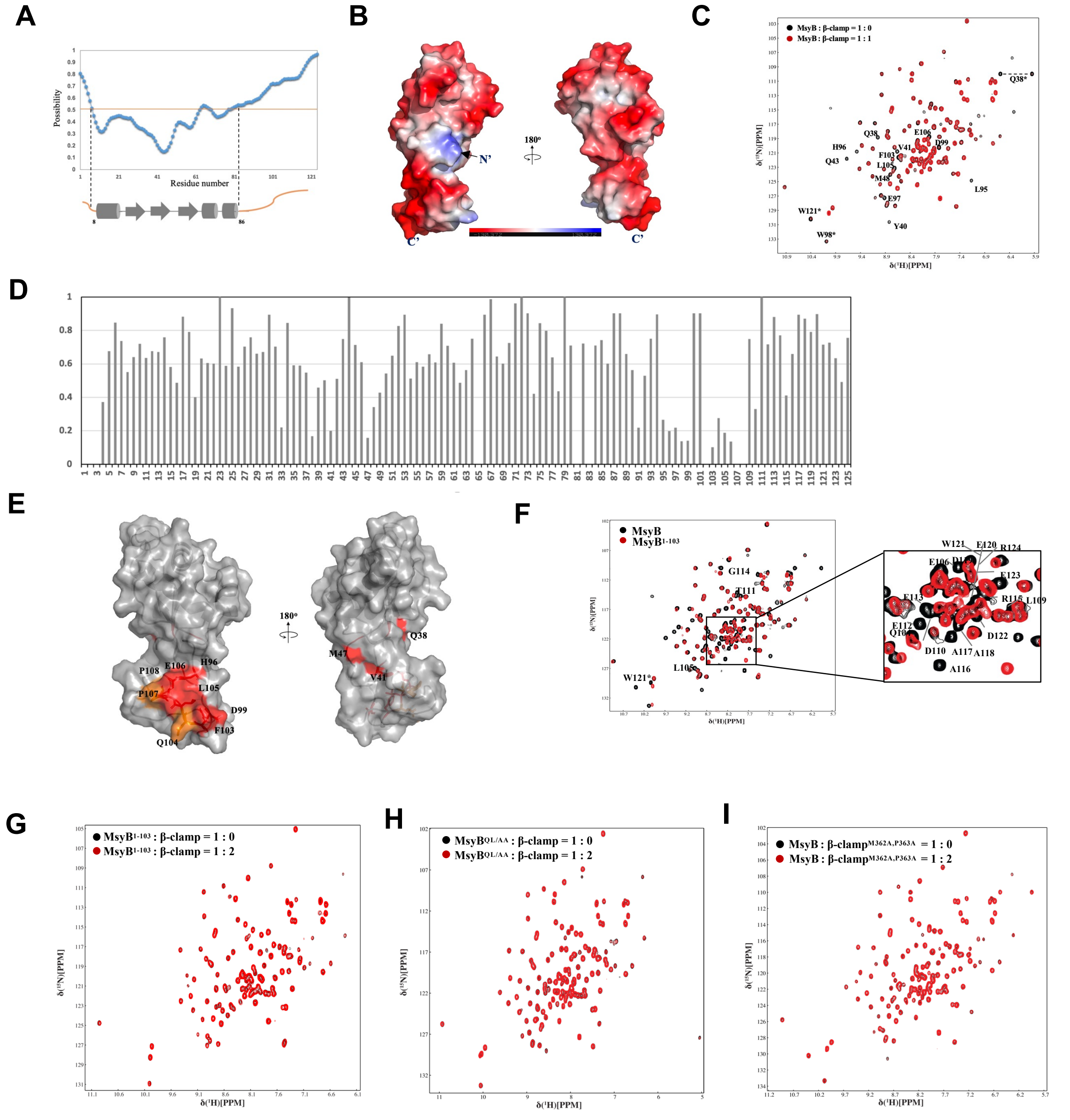

**Supplementary figure 7: MsyB and β-clamp interactions**

(**A**) The secondary structure arraignment of MsyB. Top: Intrinsically disordered protein profile predicted by IUPred3; Bottom: the topology of the solution structure. (**B**) The electrostatic surface of MsyB displayed in 180° rotation. Structure shown represents the closest structure to the average calculated by NMR. **(C)** Overlay of $^{15}$N-HSQC spectra of MsyB (black), MsyB with 1 molar equivalent of clamp added (red). **(D)** Graph of intensity ratio against residue number, showing the peak broadening effect during titration as shown in **(C)** . * indicates a proline residue while gap indicates the peak intensity which could not be extracted due to overlapping with neighbouring peaks. **(E)** The residues experiencing notable peak broadening effect (intensity ratio dropped below 20%) as shown in (**D**) are highlighted in red on the surface of MsyB with 180° rotation. Q104, P107 and P108, which are not assigned but adjacent to highlighted region, are colored in orange. **(F)** Overlay of $^{15}$N HSQC spectrum of wildtype (Black) and MsyB$^{1-103}$(Red), most residues from 104 to 125 are labelled. **(G-I)** NMR titrations show no interaction between mutants of MsyB and β-clamp as indicated.

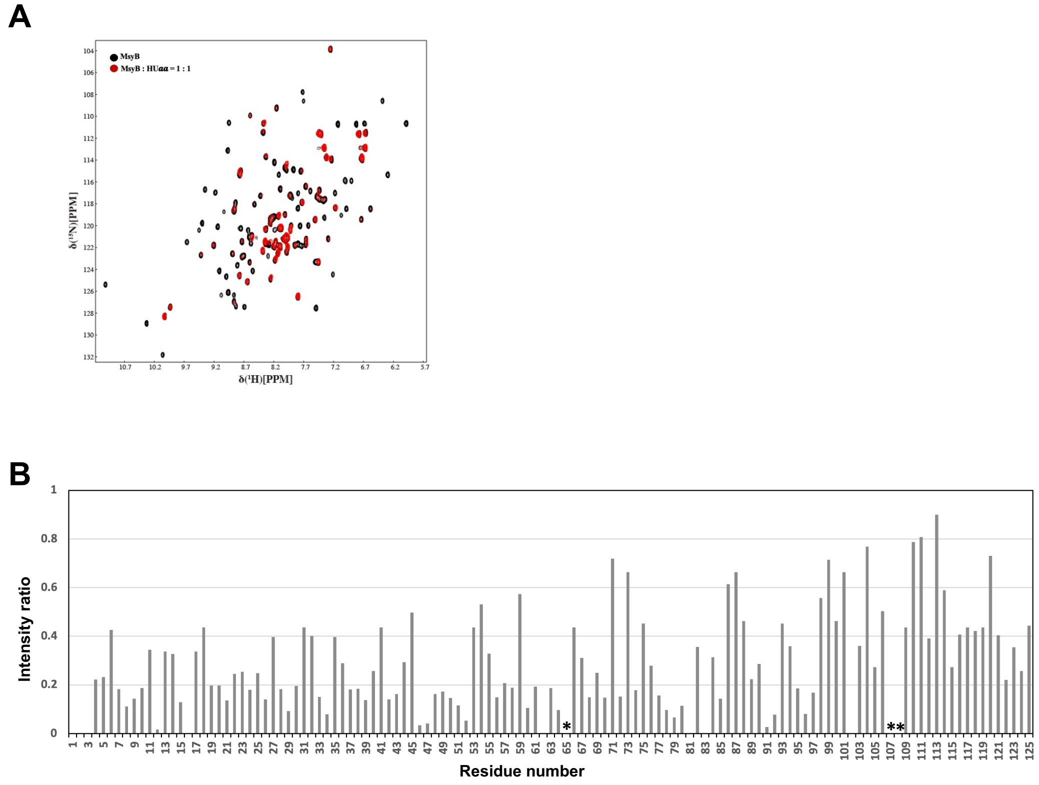

**Supplementary figure 8: MsyB and HU interactions**

(**A**) Overlay of 15N-HSQC spectra of MsyB (black) and MsyB with 1 molar equivalent of HU***aa*** of different bacteria added (Red). (**B**) Graph of intensity ratio against residue number, showing the peak broadening effect during titration as shown in **(A)**. * represents proline, and gap stands for no assignment or unable to trace due to peak overlapping. (**C**) The residues experiencing notable peak broadening effect (ratio < 0.2) as shown in (**B**) are colored in red on the surface of MsyB with 180° rotation. P65 is colored in orange.

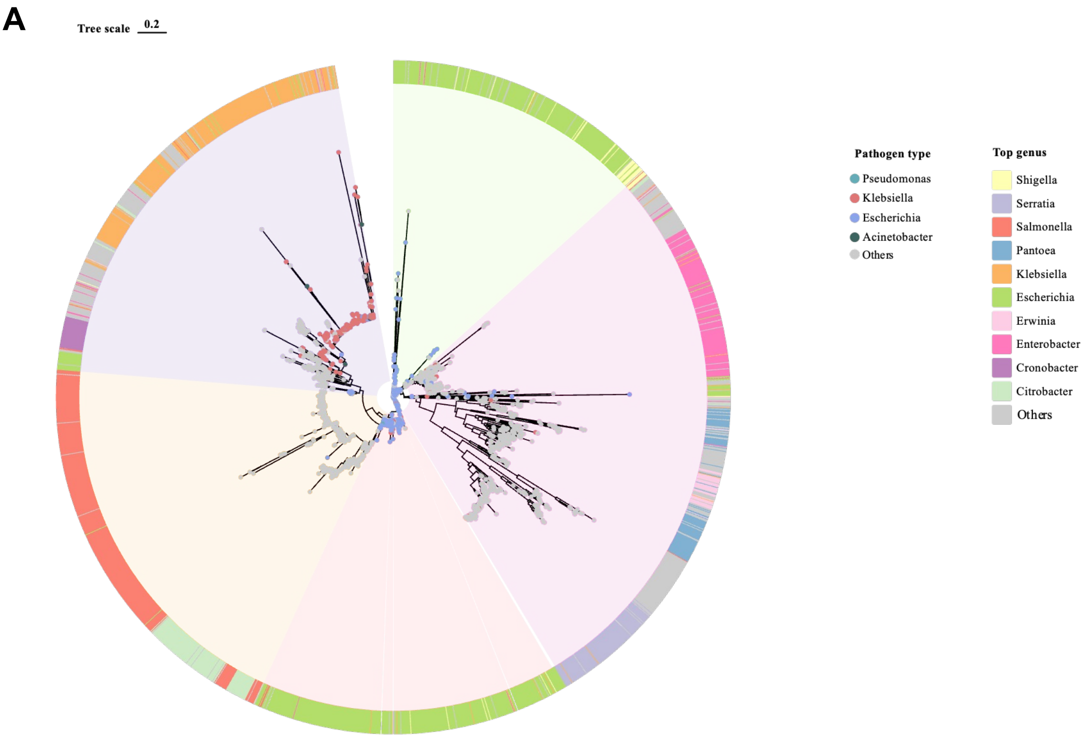

**Supplementary figure 9: The phylogenic tree of MsyB.**
The phylogenic of MsyB showing its conservation among Gram-negative bacteria, including pathogenic bacteria Pseudomonas, Klebsiella, Escherichia, Acinetobacter and others.

**Table S1. Primers used in Cloning and Mutagenesis**

| | | |
|---|---|---|
| **Plasmid** | **pET-28a-His$_6$-SUMO-MsyB** | |
| Primer | Vector Linearization | Sense: ACCACCAATCTGTTC |
| | | Antisense: TAATAGCTCGAGCAC |
| | MsyB Amplification | Sense: GAACAGATTGGTGGTATGACCATGTACGCAAC |
| | | Antisense: GTGGTGCTCGAGCTATTAACGTTCATCCCACTCATC |
| **Plasmid** | **pET-28a-His$_6$-SUMO-MsyB$^{1\text{-}103}$** | |
| Primer | MsyB$^{1\text{-}103}$ Amplification | Sense: GAACAGATTGGTGGTATGACCATGTACGCAAC |
| | | Antisense: GTGGTGCTCGAGCTATTAAAATTCCCCCTCGTCC |
| **Plasmid** | **pET-28a-His$_6$-SUMO-MsyB $^{Q104A\ L105A}$** | |
| Primer | MsyB_Q104A L105A | Sense: GGGGGAATTTGCCGCCGAGCCACCGC |
| | | Antisense: TCGTCCCATTCATGTAATG |
| **Plasmid** | **pET-28a-SUMO-MsyB-His$_6$** | |
| Primer | Vector Linearization | Sense: GGTATATCTCCTTCTTAAAG |
| | | Antisense: GGAGCAGGGGC |
| | SUMO Amplification | Sense: CTTTAAGAAGGAGATATACCATGAGCAGCGGCCTGG |
| | | Antisense: TTGCGTACATGGTCATACCACCAATCTGTTCTCTGTG |
| | MsyB Amplification | Sense: AGAGAACAGATTGGTGGTATGACCATGTACGCAACGC |
| | | Antisense: CCCAGCCCCTGCTCCACGTTCATCCCACTCATCAG |
| **Plasmid** | **pET-46-His$_6$-*E. coli* β-clamp** | |
| Primer | Vector Linearization | Sense: TAACCGGGCTTCTCC |
| | | Antisense: CTTGTCGTCGTCATCC |
| | *E. coli* β-clamp Amplification | Sense: GATGACGACGACAAGATGAAATTTACCGTAGAACGTG |
| | | Antisense: TGAGGAGAAGCCCGGTTACAGTCTCATTGGCATGAC |
| **Plasmid** | **pET-46-His$_6$-*E. coli* β-clamp$^{M362A\ P363A}$** | |
| Primer | *E. coli* β-clamp$^{M362A\ P363A}$ | Sense: atggcaCTGTAACCGGGCTTCTCC |
| | | Antisense: tgcagcGACAACATAAGCCGCGCT |

<table>
<tr><td>Plasmid</td><td colspan="2">pET46-His$_6$-B. subtilis β-clamp</td></tr>
<tr><td rowspan="2">Primer</td><td rowspan="2">B. subtilis β-clamp Amplification</td><td>Sense: GATGACGACGACAAGATGAAATTCACGATTCAAAAAGATC</td></tr>
<tr><td>Antisense: TGAGGAGAAGCCCGGTTAATAGGTTCTGACAGGAAGG</td></tr>
<tr><td>Plasmid</td><td colspan="2">pET-46-His$_6$-S. aureus β-clamp</td></tr>
<tr><td rowspan="2">Primer</td><td rowspan="2">S. aureus β-clamp Amplification</td><td>Sense: GATGACGACGACAAGATGGAATTCACTATTAAAAGAG</td></tr>
<tr><td>Antisense: GAAGCCCGGTTAGTAAGTTCTGATTGGTAAAATTAATTG</td></tr>
<tr><td>Plasmid</td><td colspan="2">pET-28a-His$_6$-SUMO-HUα</td></tr>
<tr><td rowspan="2">Primer</td><td rowspan="2">HUα Amplification</td><td>Sense: GAACAGATTGGTGGTATGAACAAGACTCAACTG</td></tr>
<tr><td>Antisense: GTGGTGCTCGAGCTATTATTACTTAACTGCGTCTTTC</td></tr>
<tr><td>Plasmid</td><td colspan="2">pET-28a-His$_6$-SUMO-HUβ</td></tr>
<tr><td rowspan="2">Primer</td><td rowspan="2">HUβ Amplification</td><td>Sense: GAACAGATTGGTGGTATGAATAAATCTCAATTGATCG</td></tr>
<tr><td>Antisense: GTGGTGCTCGAGCTATTATTAGTTTACCGCGTCTTTC</td></tr>
<tr><td>Plasmid</td><td colspan="2">pET-Duet-His$_6$-IHFα-IHFβ</td></tr>
<tr><td rowspan="8">Primer</td><td rowspan="4">Vector Linearization</td><td>Sense 1: TTAACCTAGGCTGCTGC</td></tr>
<tr><td>Antisense 1: GGTATATCTCCTTCTTAAAGTTAAAC</td></tr>
<tr><td>Sense 2: TGCTTAAGTCGAACAGAAAG</td></tr>
<tr><td>Antisense 2: ATGTATATCTCCTTCTTATACTTAAC</td></tr>
<tr><td rowspan="2">IHFα Amplification</td><td>Sense: GTTTAACTTTAAGAAGGAGATATACCATGCATCATCATCATCATCAC GGCGAGAATCTTTATTTTCAGGGCATGGCGCTTACAAAAGCTG</td></tr>
<tr><td>Antisense: ACTTTCTGTTCGACTTAAGCATTACTCGTCTTTGGGCGAAG</td></tr>
<tr><td rowspan="2">IHFβ Amplification</td><td>Sense: GTATAAGAAGGAGATATACATATGACCAAGTCAGAATTGATAG</td></tr>
<tr><td>Antisense: GGTGGCAGCAGCCTAGGTTAATTAACCGTAAATATTGGCGCG</td></tr>
</table>

| | | |
|---|---|---|
| **Plasmid** | **pET-28a-His$_6$-SUMO-Fis** | |
| Primer | Fis Amplification | Sense: GAACAGATTGGTGGTATGTTCGAACAACGCGTAAATTC |
| | | Antisense: GTGGTGCTCGAGCTATTATTAGTTCATGCCGTATTTTTTCAATTTTTTACGC |
| **Plasmid** | **pET-28a-His$_6$-SUMO-H-NS** | |
| Primer | H-NS Amplification | Sense: GAACAGATTGGTGGTATGAGCGAAGCACTTAAAATTCTG |
| | | Antisense: GTGGTGCTCGAGCTATTATTATTGCTTGATCAGGAAATCGTCGAGG |
| **Plasmid** | **pBAD-MsyB** | |
| Primer | Vector Linearization | Sense: AGGAGGAATTCACCATGG |
| | | Antisense: GCTAGCCCAAAAAAACGG |
| | MsyB Amplification | Sense: TTTTTTTGGGCTAGCATGACCATGTACGCAACGC |
| | | Antisense: TGGTGAATTCCTCCTTTAACGTTCATCCCACTCATC |
| **Plasmid** | **pTargetF-*msyB*** | |
| Primer | Vector Linearization | Sense: ACTAGTATTATACCTAGGAC |
| | | Antisense: GTTTTAGAGCTAGAAATAGC |
| | N20 Amplification | Sense: GTATAATACTAGTGAATTTCTTGCAGACAACCCGTTTTAGAG |
| | | Antisense: CTCTAAAACGGGTTGTCTGCAAGAAATTCACTAGTATTATAC |
| ***msyB*-up** | | |
| Sense | | TGTTGATCCCAATGTCTTAC |
| Antisense | | TCCTTCCTCGGTATCCAGCGTGCAGCGTCAATGGCTT |
| ***msyB*-down** | | |
| Sense | | AAGCCATTGACGCTGCACGCTGGATACCGAGGAAGGA |
| Antisense | | CACTGGCATTGCGTAAACC |

| | | |
|---|---|---|
| **Plasmid** | **pTargetF-*hupA*** | |
| Primer | N20 Amplification | **Sense:** GTATAATACTAGT GTACTGGCCGCAACCCGCAGGTTTTAG |
| | | **Antisense:** TAAAACCTGCGGGTTGCGGCCAGTACACTAGTATTATAC |
| ***hupA*-up** | | |
| Sense | | CAATCGATGCCTGCGTGGCG |
| Antisense | | CACTGCCACGCAATCAAGTTATCCTTACAATGTGTTTATC |
| ***hupA*-down** | | |
| Sense | | GATTGCGTGGCAGTGAACAGTTTTAAC |
| Antisense | | TGCCAGCGCCCCTGGC |
| **Plasmid** | **pTargetF-*hupB*** | |
| Primer | N20 Amplification | **Sense:** GTATAATACTAGT GTACTGGCCGCAACCCGCAGGTTTTAGAG |
| | | **Antisense:** CTCTAAAACCTGCGGGTTGCGGCCAGTAC ACTAGTATTATAC |
| ***hupB*-up** | | |
| Sense | | GGTTTCTTGCCTGACCGG |
| Antisense | | CTGGGGACAACGCTCTTCTCTTCCTCTTTATAATTT |
| ***hupB* -down** | | |
| Sense | | GCGTTGTCCCCAGTGGGG |
| Antisense | | CTGATCCAGCAGCGCCTCGTCGATCAG |
| **dsDNA** | | |
| DNA1 | | TGCTTATCAATTTGTTGCACC |
| DNA2 | | GGTGCAACAAATTGATAAGCA |
| **RT-qPCT** | | |
| 16S ribosomal RNA | | Sense: TGCAAGTCGAACGGTAACAGGAAG |
| | | Antisense: GGCACATCCGATGGCAAGAGG |
| *msyB* | | Sense: CCATGTACGCAACGCTTGAAGAAG |
| | | Antisense: CCACATGATGTCGCCGTCCTG |

| **Table S2: Cryo-EM structure statistics** | |
|---|---|
| **Data collection and processing** | |
| Magnification | 50,000 × |
| Voltage (kV) | 300 |
| Electron exposure ($e^-/Å^2$) | 40 |
| Defocus size (μm) | -0.5 to -2.5 |
| Pixel size (Å) | 0.95 |
| Initial particle images (no.) | 7,147,545 |
| Final particle images (no.) | 156,941 |
| Map resolution (Å) | 4.15 |
| FSC threshold | 0.143 |
| **Refinement** | |
| Model resolution (Å) | 4.2 |
| FSC threshold | 0.143 |
| Map sharpening *B* factor ($Å^2$) | -233.0 |
| Model composition | |
| Non-hydrogen atoms | 5739 |
| Protein residues | 740 |
| *B* factor ($Å^2$) | |
| Protein | min/max/mean:142.14/222.41/173.84 |
| Ligand | 0 |
| R.m.s. deviations | |
| Bond lengths (Å) | 0.004 |
| Bond angles (°) | 0.615 |
| Validation | |
| MolProbity score | 2.23 |
| Clashscore | 15.64 |
| Poor rotamers (%) | 0.00 |
| Ramachandran plot | |
| Favored (%) | 90.57 |
| Allowed (%) | 9.15 |
| Disallowed (%) | 0.27 |

### Table S3: NMR structure statistics

| MsyB | Value |
|---|---|
| **NMR Distance and Dihedral Constraints** | |
| **Distance constraints** | |
| **Total NOE** | 1718 |
| **Intraresidue** | 487 |
| **Interresidue** | 1231 |
| **Sequential (\|i-j\|)=1)** | 399 |
| **Short range (2≤\|i-j\|)≤3)** | 220 |
| **Medium range (4≤\|i-j\|)≤5)** | 85 |
| **Long range (\|i-j\|)>5)** | 527 |
| **Total Dihedral angle Restraints** | 193 |
| **φ** | 97 |
| **ψ** | 96 |
| **Total RDCs** | 0 |
| **Structural Statistics** | |
| **Violations (mean and SD)** | |
| **Distance constraints (Å)** | 0.028 ± 0.003 |
| **Dihedral angle constraints (°)** | 0.94 ± 0.091 |
| **Maximum dihedral angle violation (°)** | 0.95 |
| **Maximum distance constraint violation (Å)** | 0.30 |
| **Deviations from idealized geometry** | |
| **Bond length (Å)** | 0.004 ± 0.000 |
| **Bond angle (°)** | 0.576 ± 0.010 |
| **Impropers (°)** | 1.904 ± 0.090 |
| **Average Pairwise rmsd[α](Å)** | |
| **Heavy** | 0.379 ± 0.061 |
| **Backbone** | 0.749 ± 0.094 |
